# Systems level analysis of time and stimuli specific signaling through PKA

**DOI:** 10.1101/2022.03.10.483795

**Authors:** Michael Plank, Nicole Carmiol, Bassam Mitri, Andrew P. Capaldi

**Affiliations:** Department of Molecular and Cellular Biology, University of Arizona, Tucson, AZ 85721; the Bio5 Institute, University of Arizona, Tucson, AZ 85721

**Keywords:** Protein kinase A, PKA, carbon source, phosphorylation, transition, phosphoproteomics, sequence motif

## Abstract

Eukaryotic cells create gradients of cAMP across space and time to regulate the cAMP dependent protein kinase (PKA) and, in turn, growth and metabolism. However, how PKA responds to different concentrations of cAMP is unclear. Here, to address this question, we examine PKA signaling in *S. cerevisiae* in different conditions, timepoints, and concentrations of the chemical inhibitor 1-NM-PP1 using phosphoproteomics. These experiments show that there are numerous proteins that are only phosphorylated when cAMP and PKA activity are at/near their maximum level, while other proteins are phosphorylated even when cAMP levels and PKA activity are low. The data also show that PKA drives cells into distinct growth states by acting on proteins with different thresholds for phosphorylation in different conditions. Analysis of the sequences surrounding the 118 PKA-dependent phosphosites suggests that the phosphorylation thresholds are set, at least in part, by the affinity of PKA for each site.

## INTRODUCTION

The cAMP dependent protein kinase (PKA) is a key regulator of cell growth and metabolism in eukaryotes.

In the simple model organism *Saccharomyces cerevisiae*, PKA is (primarily) activated by glucose and other fermentable sugars via the G-protein coupled receptor Gpr1, the associated Ga protein Gpa2, and the small GTPases Ras1/2 (Broek et al., 1985; Cannon and Tatchell, 1987; Conrad et al., 2014; Kraakman et al., 1999; Rolland et al., 2000; Toda et al., 1985). Specifically, Gpa2 and/or Ras1/2 activate the adenylyl cyclase, Cyr1, to drive production of the second messenger cyclic AMP (cAMP). cAMP then binds to the regulatory subunit of the PKA holoenzyme, Bcy1, triggering the activation of the three catalytic subunits, Tpk1, Tpk2 and Tpk3 (Hixson and Krebs, 1980; Matsumoto et al., 1982; Toda et al., 1987). Tpk1-3, in turn, then phosphorylate numerous proteins to (i) prevent entry into quiescence; (ii) inhibit sporulation, autophagy, and the general stress response; (iii) activate glycolysis; and (iv) activate the phosphodiesterases Pde1/2, and other negative regulators of the PKA pathway, to reduce the concentration of cAMP over time (Budovskaya et al., 2005; Dihazi et al., 2003; Görner et al., 2002; Honigberg and Purnapatre, 2003; Hu et al., 2010; Ma et al., 1999; Munder and Küntzel, 1989; Portela et al., 2006; Reinders et al., 1998; Smith et al., 1998; Stephan et al., 2009).

In humans, and other mammals, the core of the PKA pathway—involving cAMP synthesis, the structure of the PKA holoenzyme, and negative feedback via the phosphodiesterases—is similar to that found in yeast, but the regulation of the pathway is more complex (Taylor et al., 1990; Torres-Quesada et al., 2017): First, the upstream activators vary in different tissues/cell types (Torres-Quesada et al., 2017). Second, adenylyl cyclases, phosphodiesterases, the PKA holoenzyme, and other proteins, are recruited to distinct cellular compartments by A-Kinase anchoring proteins (AKAPs) so that the pathway can drive different responses in different conditions by activating specific, localized, centers of cAMP production (Taylor et al., 1990; Torres-Quesada et al., 2017; Wong and Scott, 2004; Zaccolo et al., 2021).

Thus, from a simple perspective, it appears that the PKA pathway evolved to create gradients of cAMP, across both space and time, to precisely regulate PKA activity. However, it is currently unclear how PKA responds to different levels of cAMP.

Work in *S. cerevisiae* has begun to shed light on the impact that different concentrations of cAMP have on PKA signaling, but the data are (at least at face value) contradictory: It is well known that cAMP levels spike when glucose is added to cells growing in a non-fermentable carbon source, and then fall back to a level near the pre-stimulus baseline over approximately 20 min (Botman et al., 2021; Ma et al., 1999; Purwin et al., 1982; Thevelein and Beullens, 1985). In line with this, our recent experiments show that PKA drives a large, transient, pulse of gene expression when cells are first exposed to glucose, and this helps to speed the transition to a rapid growth state (Kunkel et al., 2019). In other words, gene expression data suggest that PKA is primarily active during the initial response to glucose, when cAMP levels are high. In contrast, studies of protein phosphorylation—virtually all of which have examined PKA signaling during steady state growth in glucose—show that PKA is active even when cAMP levels are low (Budovskaya et al., 2005; Plank et al., 2020; Searle et al., 2004; Yu et al., 2002).

One model, however, that could explain both sets of observations, is that PKA phosphorylates different proteins in different conditions/concentrations of cAMP.

Here, to test this hypothesis, we measure PKA-dependent phosphorylation across the proteome at different time-points after cells are exposed to glucose, and in different levels of the inhibitor 1-NM-PP1. These experiments reveal that, as predicted, there are numerous proteins that are only phosphorylated when cAMP levels and PKA activity are at/near their maximum level, while other proteins are phosphorylated even when cAMP levels and PKA activity are low. In fact, we find that PKA activity towards some proteins is so strong that they are phosphorylated during growth in the non-fermentable carbon source glycerol, and that PKA activity is required for cell proliferation in this condition. In addition, by analyzing the 118 PKA-dependent phosphosites identified in our study, we show that sites with a low threshold for phosphorylation tend to have a high-affinity RRXS PKA consensus motif, while those with a high threshold for phosphorylation tend to have a non-consensus, low-affinity motif, such as RXXS (Kemp et al., 1977; Shabb, 2001; Mok et al., 2010).

In sum, our data show that PKA drives the phosphorylation of different proteins in different conditions to tune the output of the pathway over time, and in response to different stimuli, and indicate that low affinity, non-consensus, phosphorylation sites play an important role in cell signaling.

## RESULTS

### PKA drives a transient response to glucose

It is well established that cAMP levels spike when glucose is added to yeast growing in a non-fermentable carbon source (such as glycerol) and then drop to a level just above the pre-stimulus baseline over a period of 10-20 min (Botman et al., 2021; Ma et al., 1999; Purwin et al., 1982; Thevelein and Beullens, 1985). Despite this, previous studies of PKA signaling have focused, almost entirely, on measuring PKA-dependent phosphorylation during steady state growth in glucose (Budovskaya et al., 2005; Plank et al., 2020; Searle et al., 2004; Yu et al., 2002). Therefore, to determine if the pulse of cAMP seen in fresh glucose activates a different signaling program than that found during log phase growth, we measured the phosphorylation changes that occur at various time-points after 2% glucose is added to cells growing in glycerol using phosphoproteomics. In the initial experiments we focused on examining signaling 0, 2.5, 5, 10, and 120 min after the cells were exposed to glucose to ensure we captured the phosphorylation changes that occur (i) when PKA activity reaches its maximum level (at 2.5, 5 and/or 10 min), and (ii) when PKA activity drops to the level found during steady state growth (120 min).

As a first step, we simply measured the phosphorylation changes that occur when cells carrying analog (1-NM-PP1) sensitive alleles of Tpk1, 2 and 3 (Tpk1-3^as^), and growing in synthetic medium with glycerol as the carbon source, are exposed to glucose (Bishop et al., 2000). In these experiments we were able to quantify the level of 18,343 peptide ions across three replicate time-courses, which were collapsed to 2596 unambiguously localized phosphosites (Table S1). The abundance of 58 of these phosphosites increased significantly (p<0.05), while the abundance of 54 of the phosphosites decreased significantly (p<0.05), and by at least 3-fold, after both 2.5 and 5 min in glucose (n=3; FDR 3.3%).

Next, to determine which, if any, of the glucose dependent phosphorylation events are driven by Tpk1-3, we carried out the same experiment, but this time treated the cells with 100nM 1-NM-PP1 [a concentration selected based on previous studies (Jorgensen et al., 2004; Kunkel et al., 2019; Lippman and Broach, 2009; Yorimitsu et al., 2007; Zaman et al., 2009)]. 77/112 glucose dependent sites had a statistically significant (p<0.05) decrease in the observed up- or downregulation found in glucose (FDR 0.4%) and were therefore defined as PKA-dependent sites (Fig. 1a). This list includes 24/54 of the sites that are down-regulated in glucose, and 50/58 sites that are up-regulated in glucose (Fig. 1a).

**Fig. 1.**
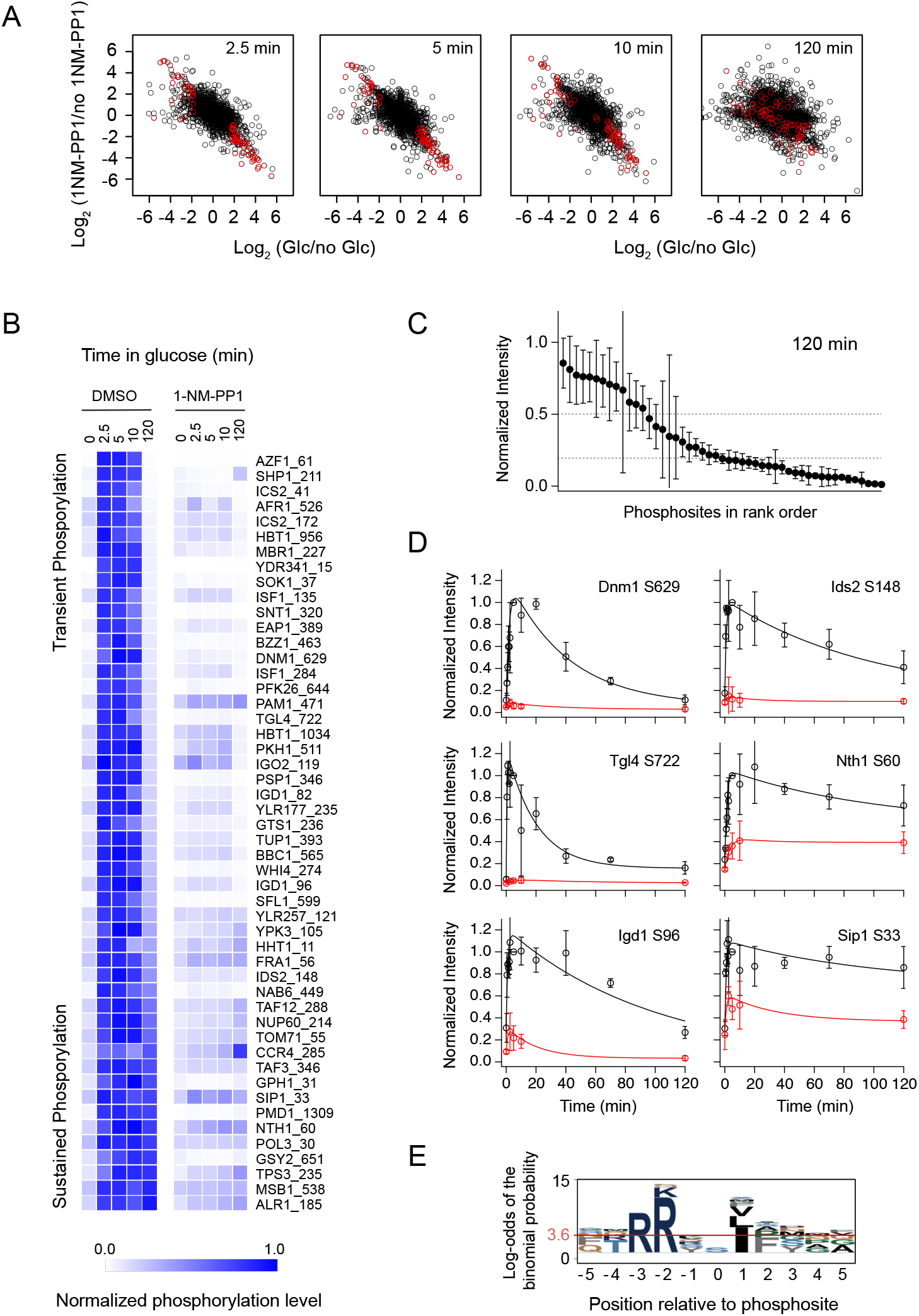
PKA-dependent protein phosphorylation during the transition from growth in glycerol, to growth in glucose. Phosphoproteomic analysis was performed to examine Tpk1-3^as^ cells harvested at various time-points after the addition of 2% glucose to cultures growing in synthetic medium with glycerol as the carbon source (SG medium). All cultures were treated with 100 nM 1-NM-PP1, or the drug carrier DMSO, 15 min prior to collection. (**a**) Changes in the abundance of 2596 phosphosites at four timepoints after glucose addition. The values shown are the mean of three replicates; the red circles highlight the PKA-dependent sites (see text for details). (**b**) Heatmap showing the phosphorylation level of the PKA-dependent sites as a function of time (normalized to the 5 min timepoint w/o 1-NM-PP1, as in all panels below). The values shown are the average from three replicates; the data from individual replicates are shown in Fig. S1. (**c**) Phosphorylation level of the PKA substrates after 120 min in glucose. The points and error bars show the average and standard deviation from three replicates. (**d**) Phosphorylation kinetics of six PKA-dependent sites after pre-treatment with 100 nM 1-NM-PP1 (red circles) or DMSO (black circles). The points and error bars show the average and standard deviation from three replicates. The lines show the best fit of each dataset to a double exponential function {Sp = A_1_*(1-exp^−k1*t^) - A_2_*(1-exp^−k2*t^) + c}. Data for other sites are shown in Fig. S2. (**e**) Sequence logo showing the amino acid sequence surrounding the PKA-dependent phosphosites identified in 100nM 1-NM-PP1.

Among the 50 phosphorylation sites that are up-regulated in glucose and have a damped response in 1-NM-PP1, most (44/50) have the characteristics of a Tpk1-3 substrate-- namely a basic residue at the −2 and/or −3 position (Table 1) (Kemp et al., 1977; Shabb, 2001; Mok et al., 2010). There is also significant enrichment for Ile, and a mild enrichment of Leu and Val at the +1 position, a known characteristic of PKA substrates in mammalian cells (Fig. 1e) (Songyang et al., 1994). It is therefore likely that most, if not all, of these 50 PKA-dependent sites are directly phosphorylated by Tpk1, Tpk2 and/or Tpk3.

**Table 1.** Motifs at the 50 PKA-dependent phosphorylation sites identified in Fig. 1. Two sites with a single lysine at the −3 position were dropped from the table since this weak motif is not enriched. The total number of sites was calculated by counting the motifs in the 2596 phosphosites quantified in this experiment. X denotes any amino acid; Z denotes any amino acid except lysine or arginine.

|  | RRXS | RKXS | KRXS | KKXS | RZXS | ZRXS | [R/K][R/K]XT |
| --- | --- | --- | --- | --- | --- | --- | --- |
| PKA dependent | 9 | 6 | 2 | 2 | 11 | 10 | 2 |
| total sites | 91 | 54 | 29 | 22 | 282 | 80 | 21 |
| enrichment | 5.1 | 5.8 | 3.6 | 4.7 | 2.0 | 6.5 | 4.9 |
| p-value | 5.4E-5 | 5.9E-4 | 0.11 | 0.07 | 0.02 | 2.6E-6 | 0.06 |

Examining the phosphorylation dynamics of the 50 glucose/PKA-dependent sites listed above revealed that there are three classes of site; 25 sites are transiently phosphorylated in response to glucose (<20% maximum phosphorylation at 120 min), 14 sites remain hyper-phosphorylated during steady state growth (>50% maximum phosphorylation at 120 min), while the remaining 11 sites have an intermediate response (Fig. 1b, c, S1). To gain further insight into the phosphorylation dynamics, we ran a separate experiment collecting samples after 0.5, 1, 1.5. 2, 2.5, 5, 20, 40, 70 and 120 min of glucose exposure. Using these data we were able to follow the precise phosphorylation dynamics of most of the 50 sites (Fig. 1d and S2) and found that (i) the PKA-dependent sites are generally phosphorylated within 1 min, and (ii) that the original classification was correct; some sites (such as Ser 60 on Nth1 and Ser 33 on Sip1) remain phosphorylated 2 hours after the transition to growth in glucose, while others (such as Ser 629 on Dnm1 and Ser 722 on Tgl4) are dephosphorylated with time constants (1/k) of 50-300 min (Fig. 1d, S2).

Overall, the time-course analysis shows that the PKA pathway does drive the phosphorylation of different substrates in different conditions; 50 sites (on 47 proteins) are phosphorylated in the immediate response to glucose (when cAMP levels are at their maximum), but only 14/50 remain highly phosphorylated during steady state growth (once cAMP levels are low). All of the 36 sites that we identified with a transient or intermediate response are novel PKA targets, including sites on: (1) the zinc finger transcription factor Azf1, known to regulate carbon metabolism and energy production in glucose; (2) Shp1 and Afr1, proteins known to regulate the PP1 phosphatase Glc7, a key regulator of glycogen metabolism, sporulation and mitotic progression; (3) Mbr1 and Isf1, paralogs thought to regulate mitochondrial function; (4) Sok1 a poorly characterized protein that, when overexpressed, rescues the growth of a Tpk1-3 deletion strain; (5) the eIF4E-associated protein Eap1 that inhibits cap-dependent translation; (6) Bzz1 and Bbc1, proteins involved in actin patch assembly/regulation; (7) Pkh1, a serine/threonine protein kinase that regulates endocytosis (Altamura et al., 1994; Bharucha et al., 2008; Böhm and Buchberger, 2013; Cosentino et al., 2000; Daignan-Fornier et al., 1994; Friant et al., 2001; Mochida et al., 2002; Newcomb et al., 2002; Slattery et al., 2006; Soulard et al., 2002; Tong et al., 2001; Ward and Garrett, 1994; Zhang et al., 1995).

### PKA drives protein phosphorylation during growth in glycerol

To our surprise, the time-course experiments described above also revealed that several of the PKA-dependent phosphosites (such as Ser 33 on Sip1 and Ser 60 on Nth1) remain partially phosphorylated in the presence of 100nM 1-NM-PP1 (Fig. 1b, d). This led us to wonder if partial inhibition of Tpk1-3 was causing us to miss some PKA substrates and/or giving us an incomplete view of PKA signaling.

To explore this possibility, we measured the growth of the Tpk1-3^as^ strain, and the base wild-type strain, in different concentrations of 1-NM-PP1. These experiments showed that 100nM 1-NM-PP1 only partly inhibits the growth of Tpk1-3^as^ cells (Fig. 2a). In contrast, 500nM 1-NM-PP1 causes near complete arrest of Tpk1-3^as^ growth, but still has no impact on wild-type cells (Fig. 2a). Therefore, to see what fraction of the PKA-dependent phosphorylation program we were missing in our first round of experiments, we measured PKA signaling 0 and 5 min after exposure to glucose, as described earlier, but this time using 500nM 1-NM-PP1. Remarkably, 500nM 1-NM-PP1 not only had a larger overall impact on protein phosphorylation, but it also caused the abundance of some (but not all) phosphosites to drop significantly below the level found during steady state growth in glycerol, as discussed in detail below. These data suggested that there is significant PKA kinase activity even in the non-fermentable carbon source glycerol, and that different PKA substrates have different sensitivities to 1-NM-PP1.

**Fig. 2.**
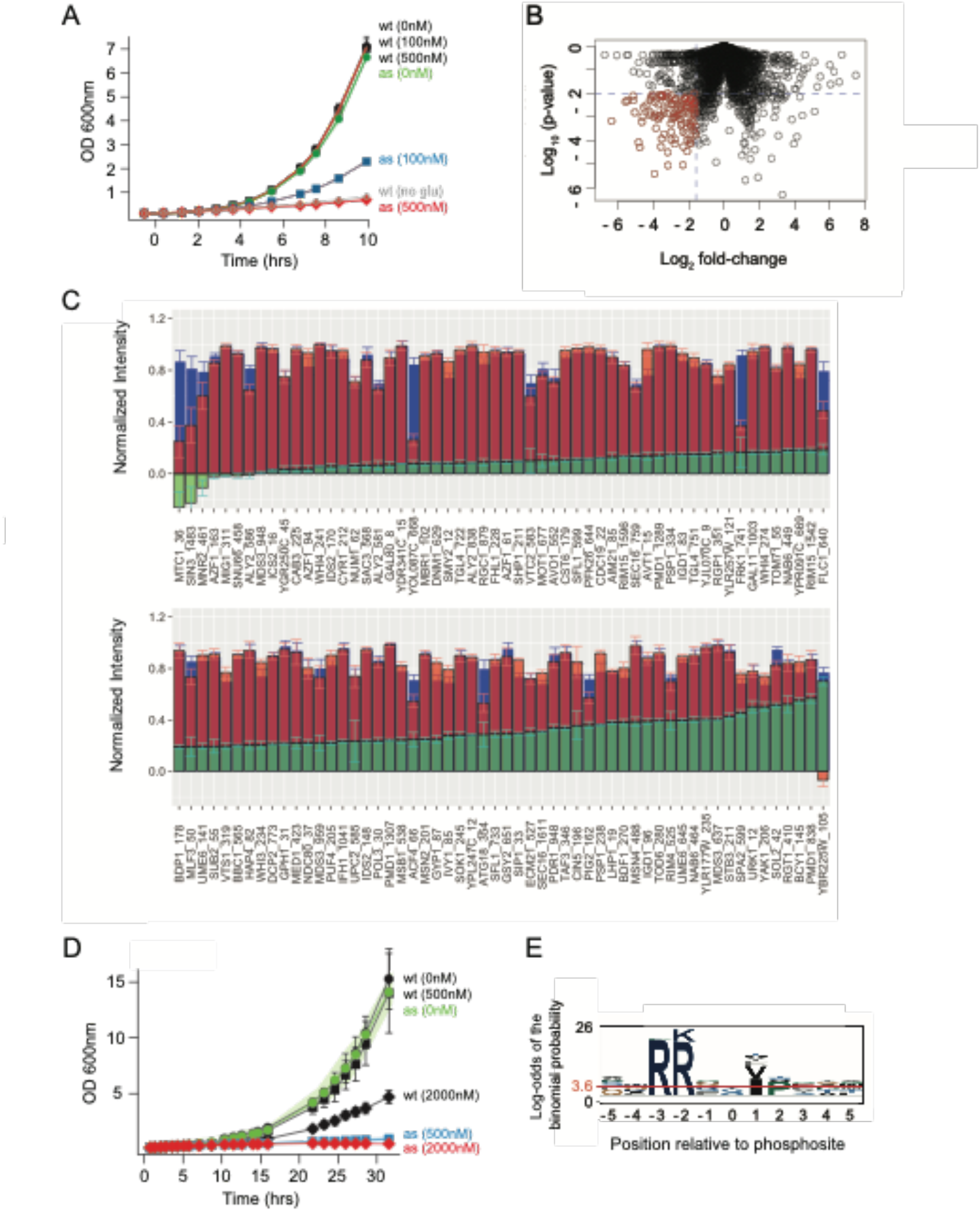
PKA-dependent protein phosphorylation before and after the addition of glucose to cultures growing in SG-medium. Phosphoproteomic analysis was performed as described in Fig. 1, except that 500 nM 1-NM-PP1 was used instead of 100 nM 1-NM-PP1. (**a)** Proliferation of Tpk1-3^as^ and wt strains after the addition of glucose (t = 0) to cultures grown in SG-medium pre-treated with the indicated concentration of 1-NM-PP1. The points and error bars show the average and standard deviation from three replicates. Note that most error bars are smaller than the points. The grey points and line show the data for cultures grown without glucose or 1-NM-PP1. Cultures were diluted once they reached OD ~0.8 and the dilution factor incorporated into the OD values shown. (**b)** Volcano plot comparing the average phosphorylation level of 4384 phosphosites after 5 min in glucose with, and without, 500 nM 1-NM-PP1 pretreatment (n=3). The dashed blue lines show the cut-offs used to identify the PKA-dependent sites (shown in red). (**c**) Conditiondependent phosphorylation at the PKA-dependent sites. **Green bars** show the level of PKA-dependent phosphorylation in glycerol, calculated by comparing the normalized phosphorylation level in 500 nM 1-NM-PP1 in SG medium, to that in 0 nM 1-NM-PP1 in SG medium. **Red bars** show the increase in phosphorylation at the PKA sites after 5 min in glucose, calculated by comparing the normalized phosphorylation level before, and 5 min after, the addition of glucose to cultures growing in SG medium. **Blue bars** show the total PKA-dependent phosphorylation at each site, calculated by comparing the normalized phosphorylation level in the cultures treated with glucose and 500nM 1-NM-PP1 to those treated with glucose and no 1-NM-PP1. All values are normalized to the phosphorylation level after 5 min in glucose without 1-NM-PP1. The error bars show the standard error of the mean, and six sites with standard errors above 0.33 are omitted from the plot. (**d**). Proliferation of Tpk1-3^as^ and wt strains in SG medium treated with the indicated concentration of 1-NM-PP1. The points and error bars show the average and standard deviation from three replicates. Note that some error bars are smaller than the points, and the error for the Tpk1-3^as^ strain is shown by a green shadow for clarity. Cultures were diluted once they reached OD ~0.8, and the dilution factor incorporated into the OD values shown. (**e**) Sequence logo showing the amino acid sequence surrounding the PKA-dependent phosphosites identified in 500nM 1-NM-PP1.

To follow up on these observations, we set up a large scale experiment where we (i) grew Tpk1-3^as^ cells in glycerol and treated them with 500nM 1-NM-PP1 or the drug carrier DMSO for 15 min and (ii) grew cells in glycerol, treated them with 0, 2.5, 5, 10, 20, 40, 80, 160 or 500nM 1-NM-PP1 for 10 min, added glucose to the cultures, and then harvested the cells after 5 min (all in at least duplicate). In these experiments, we were able to quantify 26,971 peptide ions across the samples, resulting in 4384 unambiguously localized phosphosites.

118 of the 4384 phosphosites had a significant (p<0.01) and >3-fold decrease in abundance due to treatment with 500nM 1-NM-PP1 in glucose (n=3, FDR=2.9%) and were therefore designated as PKA-dependent phosphosites (Fig. 2b). As expected, the majority (83/118) of these sites have the characteristics of a Tpk1-3 substrate, including basic residues at position −2 and/or −3, and significant enrichment for Ile and Val at the +1 position, and are therefore probably direct Tpk1, Tpk2 and/or Tpk3 targets (Table 2, Fig. 2e)

**Table 2.** Motifs at the 118 PKA-dependent phosphorylation sites identified in Fig. 2. Three sites with a lysine at the −3 position, and one site with a lysine at the −2 position, were dropped from the table since these weak motifs are not enriched. The total number of sites was calculated by counting the motifs in the 4384 phosphosites quantified in this experiment. X denotes any amino acid; Z denotes any amino acid except lysine or arginine.

|  | RRXS | RKXS | KRXS | KKXS | RZXS | ZRXS | [R/K][R/K]XT |
| --- | --- | --- | --- | --- | --- | --- | --- |
| PKA dependent | 28 | 14 | 6 | 4 | 20 | 11 | 4 |
| total sites | 123 | 76 | 74 | 41 | 446 | 147 | 42 |
| enrichment | 8.5 | 6.8 | 3.0 | 43.6 | 1.7 | 2.8 | 3.5 |
| p-value | 3.1E-18 | 2.1E-8 | 0.02 | 0.03 | 0.02 | 2.0E-3 | 0.03 |

Comparing the impact that 500nM 1-NM-PP1 has on cells growing in glycerol to the impact it has on cells in glucose, confirmed our initial finding; PKA drives protein phosphorylation in glycerol (green bars, Fig. 2c) but then becomes hyper-activated upon exposure to glucose (red bars, Fig. 2c). The data also showed that the level of PKA-dependent phosphorylation in glycerol varies dramatically from substrate to substrate: Proteins like Yak1, Stb3, Tod6, and Msn2/4 (involved in cell growth control and stress responses), and Pmd1, Ume6, Mds3, Rim4 and Bdf1 (involved in the regulation of meiosis and sporulation) are heavily phosphorylated in glycerol and have <2-fold increase in phosphorylation during the immediate response to glucose (Fig. 2c) (Broach, 2012; Deng and Saunders, 2001; García-Oliver et al., 2017; Garrett et al., 1991; Görner et al., 1998; Huber et al., 2011; Jorgensen et al., 2004; Neiman, 2011). In contrast, proteins like Mig1 and Azf1 (transcription factors involved in glucose signaling), Num1 and Dnm1 (proteins involved in mitochondrial organization), and Snu66 and Sac3 (proteins involved in ribosome biogenesis), are weakly phosphorylated in glycerol, and have a >5-fold increase in phosphorylation upon exposure to glucose (Fig. 2c) (Nehlin and Ronne, 1990; Otsuga et al., 1998; Slattery et al., 2006; Klecker et al., 2013; Stevens and Abelson, 1999; Li et al., 2009).

In line with these observations, we found that PKA activity is required for cell growth in synthetic medium with glycerol as the carbon source (Fig. 2d), and not just growth in medium containing glucose (Fig. 2a).

### Individual proteins have different sensitivities to PKA

The finding that some PKA targets are phosphorylated when cAMP levels are low (in glycerol), while others are only phosphorylated when cAMP levels are high (after 5 min in glucose), suggests that (i) different PKA substrates have different sensitivities to Tpk1-3 activity, or (ii) changes in phosphatase activity, protein localization and/or other factors modulate the output of the PKA pathway as cells switch from growth in glycerol, to growth glucose. To distinguish between these models, we turned to the 1-NM-PP1 titration experiment described above. These data show that the Tpk1-3 target sites have a wide range of sensitivities to PKA activity (Fig. 3a), consistent with model (i).

**Fig. 3.**
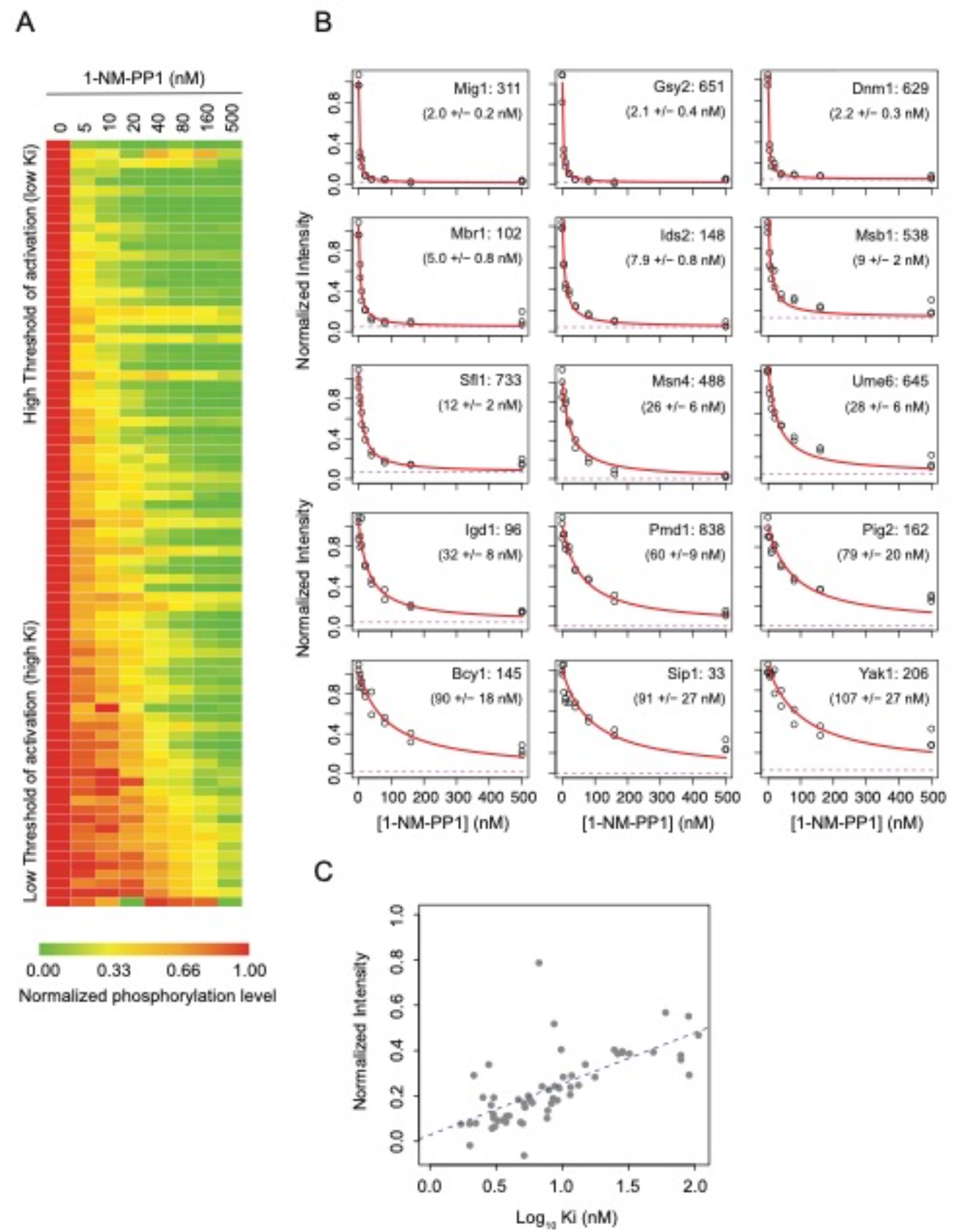
Sensitivity of PKA-dependent phosphosites to PKA inhibition. Tpk1-3^as^ cells were grown in SG medium, treated with the indicated concentration of 1-NM-PP1 for 10 min, and then exposed to glucose for 5 min before analysis using phosphoproteomics (**a**) Heatmap showing the dose-response of PKA-dependent sites (from Fig. 2b) that contain at least one arginine or two lysines in the P-3 and P-2 positions. The map shows the average value of two or three replicates and all signals are normalized to the maximum at the given phosphosite. (**b**) 1-NM-PP1 dose-response curves for 15 representative phosphosites (data for all others phosphosites are shown in Fig. S4). The red curves show the best fit to the a standard binding equation (K_i_/(K_i_ +[NM-PP1]) + c; where K_i_ is the inhibition constant and c is the baseline level of phosphorylation). The dashed violet lines show the best fit value of c. (**c)**Correlation between the K_i_ value for each phosphosite and its 1-NM-PP1-dependent phosphorylation level in SG medium (green bars Fig. 2c).

To analyze the 1-NM-PP1 titration data in more detail, we fit the data for each of the 118 PKA-dependent phosphosites to a standard inhibition curve (Fraction phosphorylated = K_i_/(K_i_ +[NM-PP1]) + c, where K_i_ is the inhibition constant and c is the baseline level of phosphorylation at a given site). This simple equation fit the data very well and showed that there is a 50-fold range in the sensitivity to 1-NM-PP1, with inhibition constants ranging from 2nM to 100nM (Fig. 3b; Fig. S4a and b).

To see if these differences explain why some proteins are phosphorylated during growth in glycerol, while others are only phosphorated during the immediate response to glucose, we examined the relationship between the K_i_ for each site and its phosphorylation level in glycerol. This analysis gave a clear answer: the sites that are highly phosphorylated in glycerol have the lowest threshold for phosphorylation by Tpk1-3 (highest K_i_ values; p-value on slope <1E-8, Fig. 3c).

Thus, the phosphoproteomic data provide strong support for a revised model of PKA signaling: PKA is driven into different activity levels in different conditions; very low in glycerol, high in the immediate response to glucose, and low/moderate during steady state growth in glucose. Each condition then triggers the phosphorylation of a subset of the PKA targets, selected in a large part, based on their sensitivity to Tpk1-3.

### Mechanisms underlying the differential sensitivity to PKA

Our discovery that different PKA targets have different sensitivities to kinase activity led to the obvious question; how do these different sensitivities arise? To address this question, we considered four possible mechanisms:

I. *The different PKA isoforms, Tpk1-3, are activated by different concentrations of cAMP and then phosphorylate different proteins*. To test this hypothesis, we built three additional strains, each carrying two out of the three analog-sensitive PKA alleles (Tpk1/2^as^, Tpk1/3^as^, Tpk2/3^as^) and used them to examine the phosphorylation levels that are achieved when a single PKA isoform is active. These experiments show that Tpk1, Tpk2 and Tpk3 act largely redundantly during growth in glycerol (Figs. 4a and S5). They also show that Tpk2 can drive (most of) the response to glucose on its own, with Tpk1 acting redundantly/partially redundantly at many sites (Fig. 4b). Thus, the PKA pathway does not rely on the activation of different PKA isoforms to create different outputs since Tpk2 can create the observed multi-level response on its own. However, the precise phosphorylation level of each protein may be tuned by its sensitivity to the individual Tpk isoforms.
II. *PKA has different affinities for different proteins, and this sets the threshold for activation at each target site*. It is difficult to calculate/estimate the strength of the interaction between a kinase and substrate since it can, in theory, be controlled by numerous factors. However, PKA is thought to bind to its substrates—in a large part— through interactions between the active site and residues near the target Ser/Thr, and is known to have a preference for the consensus RRXS motif, followed by RKXS, KRXS and then RXXS and XRXS motifs (Mok et al., 2010). We therefore examined our data to see if it is the high-affinity consensus sites that are phosphorylated in glycerol. This did turn out to be the case; 24/34 of the sites with a consensus RRXS motif have above the median level of phosphorylation in glycerol (p<0.002, hypergeometric test). In line with this, sites with an RRXS motif tend to be phosphorylated at a higher level in glycerol than sites with other, weaker motifs (p<0.001, Benjamini-Hochberg corrected value from a Mann-Whitney test against the combination of all other groups; Fig 5a).
III. *Different PKA substrates have different subcellular localizations, increasing or decreasing their access to active PKA, thus altering their threshold for activation*. Studies of Tpk1-3 localization in yeast have shown that the kinases move between the nucleus, cytoplasm, and stress granules (Tudisca et al., 2010). We therefore classified each of the PKA substrates according to its subcellular localization using the LoQAtE database and, in a few cases, primary literature (Breker et al., 2014). Very few substrates were located outside of the nucleus or cytoplasm, and thus we scored the substrates as nuclear, cytoplasmic, or unknown/other. Comparing these classifiers to our phosphoproteomic data, revealed that the substrates with nuclear localization tend to be phosphorylated at a higher-level during growth in glycerol, with 24/33 nuclear substrates having greater than the median level of phosphorylation (p<0.001, hypergeometric test). This trend also holds when looking at the localization of substrates with specific motifs (Fig. 5a).
IV. *The substrates that are heavily phosphorylated in glycerol are dephosphorylated slowly, to lower the threshold for activation by PKA*. The phosphorylation level of a given site is determined by the ratio of its phosphorylation and dephosphorylation rates. Thus, the phosphorylation levels in glycerol may be set, at least in part, by the relevant phosphatase(s). To see if this is the case, we exposed Tpk1-3^as^ cells growing in glycerol to glucose for 5 min (to drive the substrates into the phosphorylated state), treated them with 500nM 1-NM-PP1, and then measured the level of phosphorylation after 0, 1, 2.5, 5, 10, 15, 30 and 60 min (all samples in duplicate; Fig. 6a and S7). Using these data, we were able to follow the dephosphosphorylation of 58 of the 118 sites described above across the whole time-course, and the data for 46 of these 58 sites fit well to an exponential decay (relative standard error <0.33; Figs. 6b and S7a). We then examined the correlation between the rate constant for the dephosphorylation (k_dep_) of each substrate and its phosphorylation level in glycerol. This analysis showed a weak trend; high affinity sites with a low level of phosphorylation in glycerol tend to have faster dephosphorylation rates (p=0.14, Fig. 5b). In sum, our analysis suggests that the affinity of each protein for the Tpk1-3 kinases, its localization, and its dephosphorylation rate, all play a role in setting its threshold for phosphorylation and allow the PKA pathway to activate different phosphorylation programs in glycerol and glucose.

**Fig. 4.**
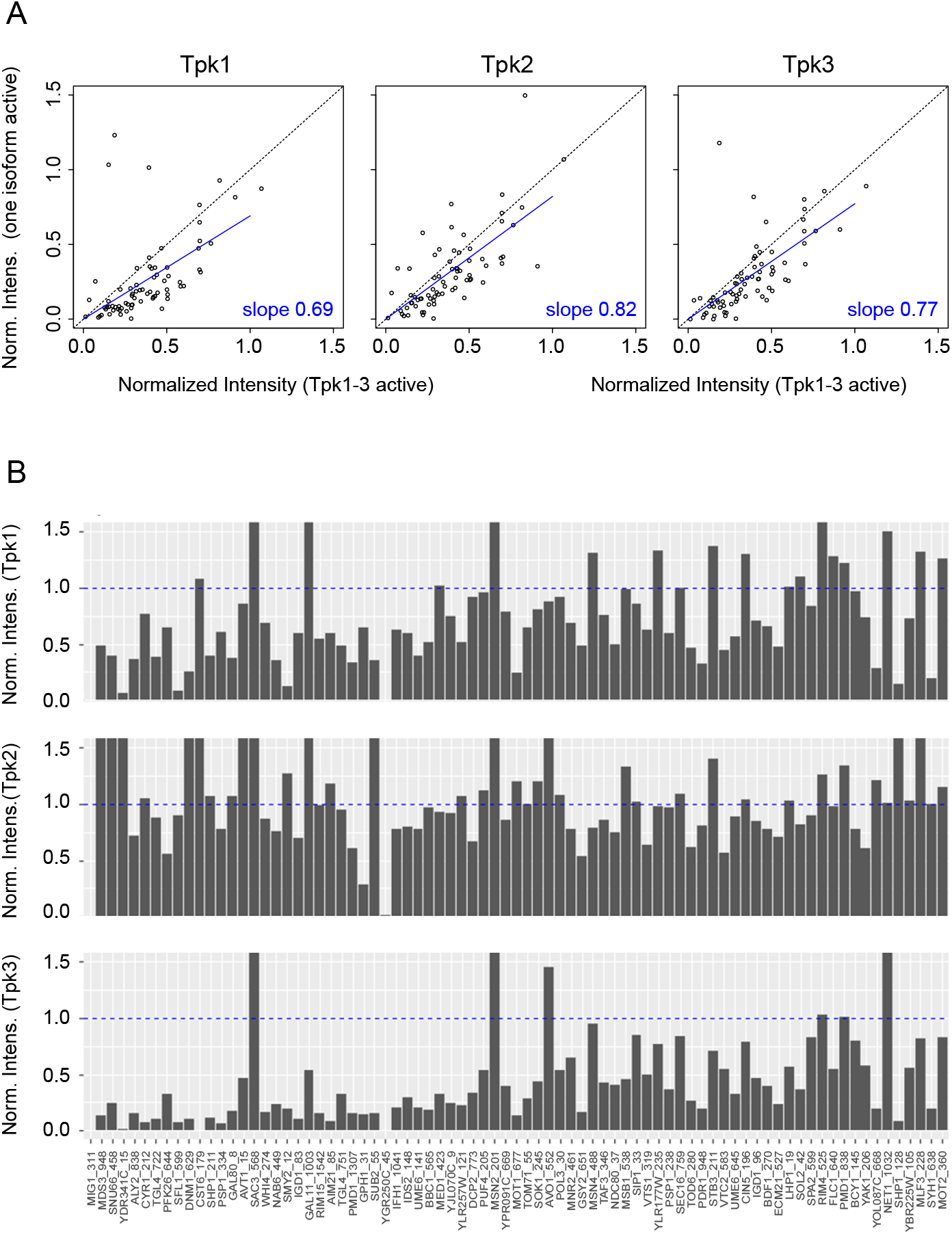
Influence of the three Tpk isoforms on protein phosphorylation. **(a)** Correlation between the phosphorylation level in SG medium in a culture with a single Tpk isoform active versus a culture with all three Tpk isoforms active. The blue lines show the best fit to a straight line with a y-intercept of zero. All values are the average from two replicates and are normalized to the phosphorylation level 5 min after glucose is added to a culture growing in SG medium. (**b**) Bar plots showing the phosphorylation level of the PKA-dependent sites 5 min after glucose is added to a culture with a single active Tpk isoform. The dashed blue lines show the phosphorylation level when all three isoforms are active. Data are shown for 79 of the 118 PKA-dependent sites identified in Fig. 2b; other sites could not be detected/quantified in this experiment.

**Fig. 5.**
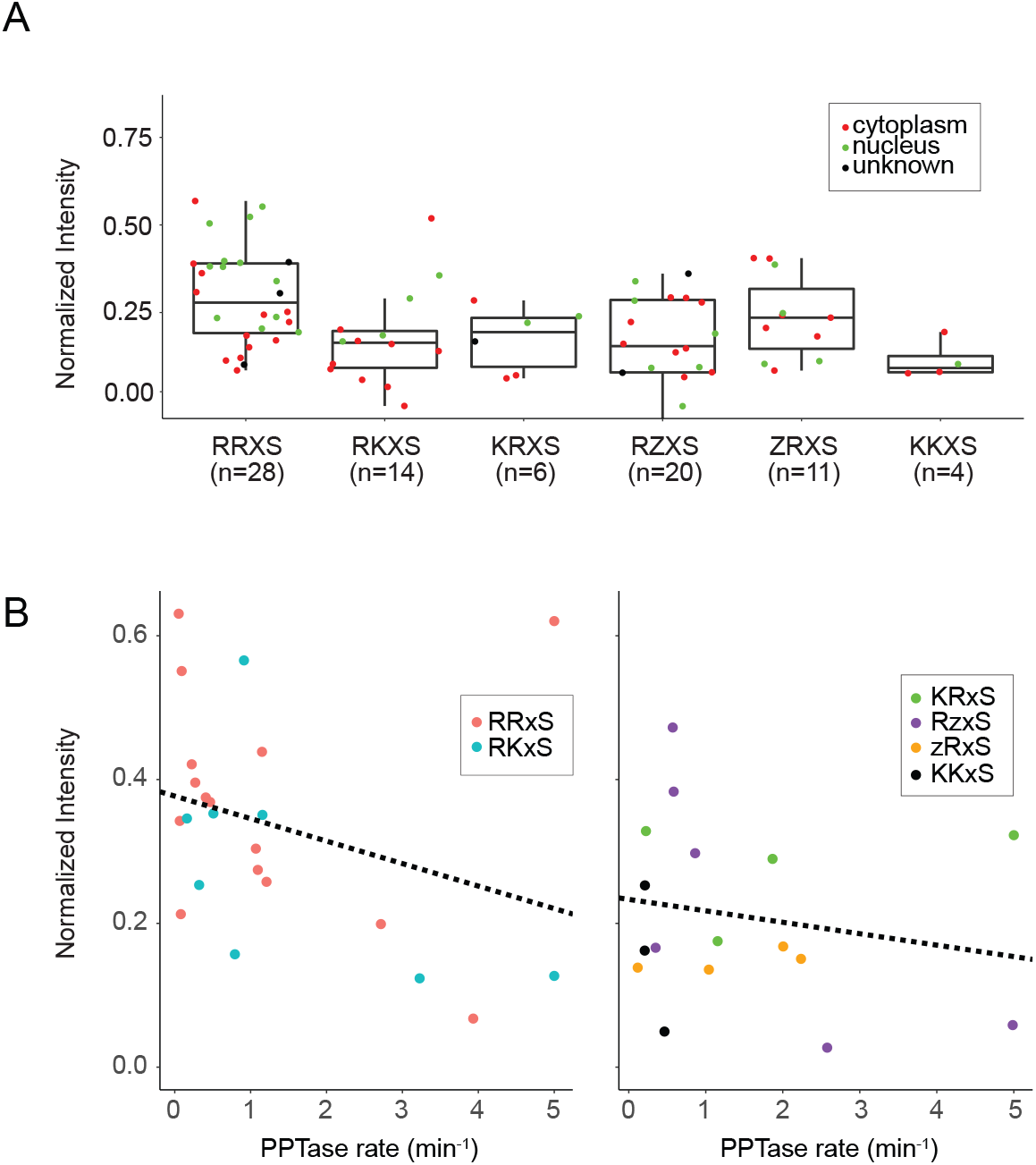
Influence of sequence, subcellular localization, and phosphatase rates on the phosphorylation levels in SG medium. (**a**) Association between the PKA-dependent phosphorylation level in glycerol (green bars, Fig. 2c) and each substrate’s sequence motif and subcellular localization. Boxes depict median and quartiles, whiskers extend to the most extreme values, but no further than 1.5× the inter-quartile range. Individual data points are color coded according to protein localization. (**b**) Correlation between the rate of dephosphorylation (from Fig. 6) and the phosphorylation level in SG, for each PKA-dependent substrate. Data were split into those for sites associated with high affinity RRXS or RKXS motifs (left panel) and those with weaker binding motifs (right panel) and fit to a linear model (black lines).

**Fig. 6.**
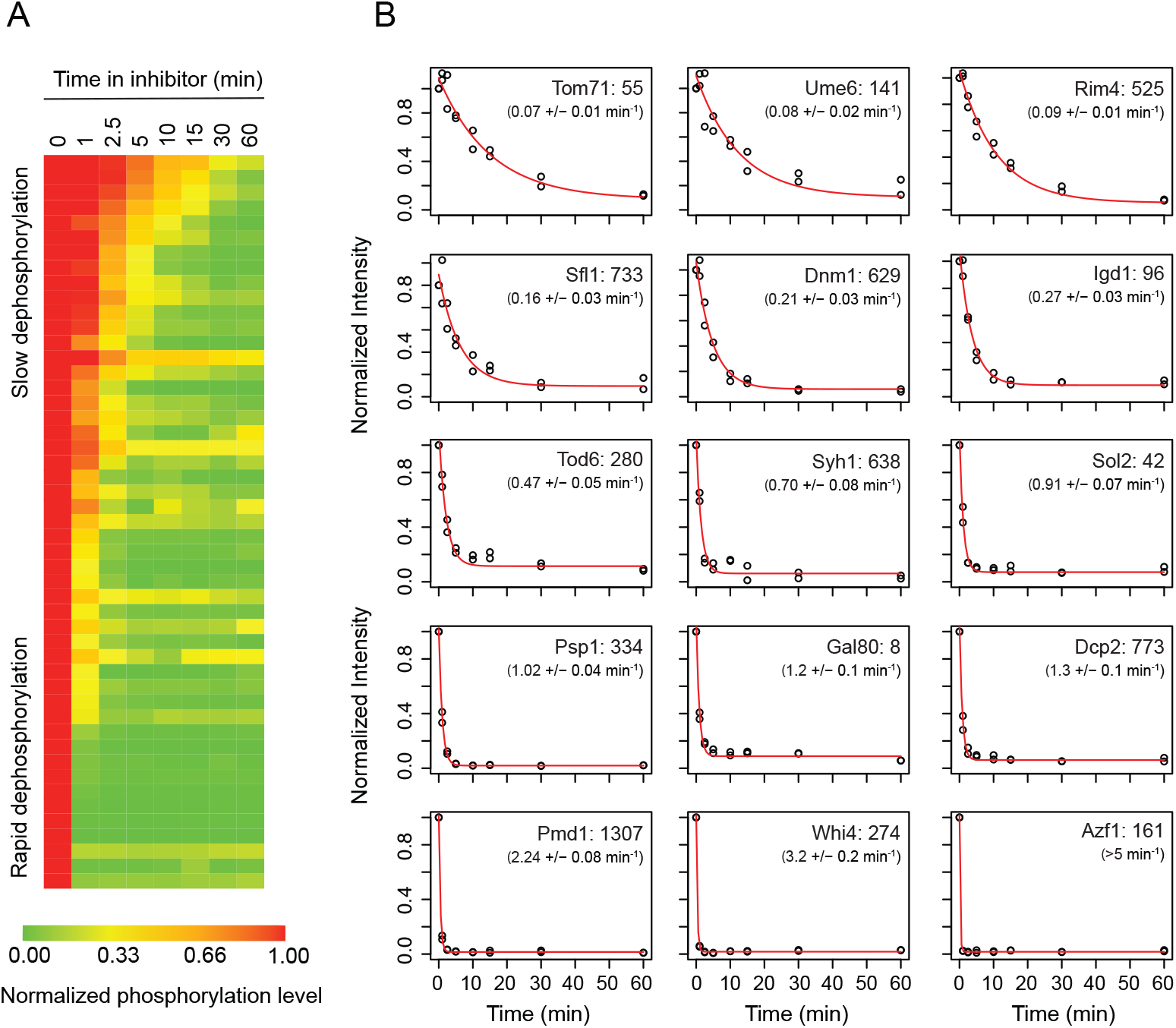
PKA-dependent phosphorylation decreases rapidly after PKA inhibition. Tpk1-3^as^ cultures were grown in SG medium, and glucose added to the culture for 5 min to drive protein phosphorylation, before 500 nM 1-NM-PP1 was added to the culture and the phosphorylation levels measured over time. (**a**) Heatmap showing the phosphorylation level of PKA-dependent sites (identified in Fig. 2b) that contain at least one arginine or two lysines in the P-3 and P-2 positions (data are the average to two replicates). Phosphorylation levels were normalized to the maximum value at each phosphosite. Note that not all PKA-dependent sites were detected/quantified in this experiment. (**b**) The phosphorylation level at various timepoints shown for 15 representative proteins (fits to all data are shown in Fig. S7). The red line shows the best fit to an exponential equation (Sp = Ae^−kt^ + c).

## DISCUSSION

Most models of cell signaling assume that kinases activate/repress their target proteins in a synchronized fashion, acting as a global switch or rheostat. Here we show that this assumption is false--at least for PKA in budding yeast. Instead, Tpk1-3 drive at least three distinct phosphorylation programs: First, when cells are growing in the non-fermentable carbon source glycerol, Tpk1-3 are active at a low level and phosphorylate approximately 30 proteins. Then, once cells are exposed to glucose, and cAMP levels spike, Tpk1 and Tpk2 drive the phosphorylation of approximately 80 additional proteins. Finally, as cells adapt to growth in glucose, and cAMP concentrations drop, the phosphorylation levels of most (but not all) of the PKA substrates decreases.

The data presented here also show that PKA drives cells into different signaling states by acting on proteins with different thresholds for phosphorylation. Specifically, we show that the sites that are highly phosphorylated in glycerol are the most sensitive to Tpk1-3, while those that are only phosphorylated during the cAMP pulse in glucose are the least sensitive to Tpk1-3. Furthermore, we show that the phosphorylation level of each site in glycerol correlates with (i) its affinity for Tpk1-3 as defined by the sequence around the target Ser/Thr, (ii) its subcellar localization, and to a lesser extent (iii) the rate of dephosphorylation.

### But why does the PKA pathway function this way?

We propose that PKA is driven to different activity levels to coordinate three distinct, but overlapping, responses: In the presence of adequate nutrients, and a non-fermentable carbon source, Tpk1-3 are activated at a low level to phosphorylate high sensitivity substrates such as Msn2/4, Yak1 and Tod6 to prevent entry into quiescence/stationary phase (Broach, 2012), and Ume6, Rim4 and Bdf1 to suppress sporulation/meiosis (Deng and Saunders, 2001; García-Oliver et al., 2017; Neiman, 2011) (Fig. 2c). In contrast, when cells are first exposed to a fermentable carbon source (and other key nutrients) PKA is fully activated for a short period of time. This triggers phosphorylation of all the PKA substrates, to push cells out of quiescence and/or sporulation (if they are not already growing with low PKA activity), but also drives a pulse of gene expression and other reprogramming events to speed the transition into a rapid growth state. Then as the cells adapt to growth in glucose, PKA activity falls to a level just above that found in non-fermentable carbon source, so that Tpk1-3 continue to phosphorylate proteins such as Tps3, Gsy2, and Nth1 involved in carbohydrate storage (François and Parrou, 2001), Sip1 involved in Snf1/AMPK regulation (Schmidt and McCartney, 2000), and Ccr4, Taf3 and Taf12 involved in Pol II function (Laribee et al., 2015; Lee and Young, 2000), to sustain fermentative growth (Fig. 1).

Given the advantages of controlling a multi-level response using a single pathway/kinase, it seems likely that other kinases will function like Tpk1-3. In fact, the Nurse lab has already shown that a single Cyclin/Cdk pair can drive the ordered phosphorylation of proteins during the cell cycle, and that this behavior depends on the fact that the target proteins have different sensitivities to CDK (Swaffer et al., 2016). Moving forward, it will therefore be important to analyze the input-output behavior of other key kinases on a proteome-wide scale to determine how they act in different types, levels and durations of stimuli. Doing so will not only give us a clearer view of the function of each kinase but should also make it possible to accurately predict the impact that mutations and drugs have on kinase function.

## MATERIALS AND METHODS

### Yeast cultures and treatment

*S. cerevisiae* strains Tpk1-3^as^ (W303; tpk1-M164G, tpk2-M147G, tpk3-M165G), Tpk1/2^as^, Tpk1/3^as^, Tpk2/3^as^, or the base wild-type strain, were inoculated into synthetic medium containing 3% glycerol as a carbon source (SG) and grown shaking at 200 rpm and 30 °C. Cultures were then diluted to a low optical density (OD_600_ ~0.02) on day 3, and again on day 4, to keep them in log phase. Once the cultures reached OD_600_ ~0.3 on day 5, 2% glucose (or an equivalent amount of water) was added to each flask using a 40% stock. Cultures were treated with the indicated amount of 1-(1,1-dimethylethyl)-3-(1-naphthalenyl)-1H-pyrazolo[3,4-d]pyrimidin-4-amine (1-NM-PP1; Millipore Sigma), taken from a 10,000x stock in DMSO, 15 min before cell harvest, unless noted otherwise. This treatment protocol was selected to ensure: (1) that we monitored the primary (and not secondary) effects of PKA inhibition and (2) that protein dephosphorylation always proceeded for the same length of time.

### Experimental design

To discover PKA substrates using 100 nM 1-NM-PP1 (Fig. 1a-c; # 4409 in tab. S1), three Tpk1-3^as^ cultures were grown until day 4, and each was split into ten 50 ml cultures. Samples were then harvested 0, 2.5, 5, 10 and 120 min after glucose addition, both with and without 100 nM 1-NM-PP1 pre-treatment.

To refine our measurement of the PKA-dependent phosphorylation kinetics (Fig. 1d and S2; #4449), two Tpk1-3^as^ cultures were grown to day 4. 50 ml of each culture was then transferred into a fresh flask, treated with 100 nM 1-NM-PP1 for 10 min, and glucose added for 5 min before harvest. Samples were then collected from the original flasks without 1-NM-PP1 addition, immediately before and 5, 10, 20, 40, 70 and 120 min after glucose addition. In a second experiment, two Tpk1-3^as^ cultures were grown to day 4, each culture was split into nine 50 ml aliquots, and cultures harvested before and 0.5, 1, 1.5, 2, 2.5 and 5 min after glucose addition (Fig. 1d and S2; #4459). The two remaining cultures from each sample were harvested 2.5 and 5 min after glucose addition with 1-NM-PP1 pre-treatment.

To discover PKA substrates using 500 nM 1-NM-PP1 and measure the sensitivity of each phosphosite to inhibitor (Fig. 2 and 3; #4520), four Tpk1-3^as^ cultures were grown to day 4. Two of the cultures were then split into ten flasks each. Nine of these flasks were pretreated with 0, 2.5, 5, 10, 20, 40, 80, 160 or 500 nM 1-NM-PP1 (in random order), and glucose added 5 min before harvest. DMSO, and H_2_O in place of glucose, was added to the remaining culture and harvested after 5 min as a control. The remaining two large scale cultures were split into four flasks each. Two were pre-treated with DMSO, and then one treated with glucose, and one treated with H_2_O, 5 min before harvest. The other two cultures were pre-treated with 500 nM 1-NM-PP1, and one treated with glucose, and the other treated with H_2_O, 5 min before harvest.

In an independent 1-NM-PP1 titration experiment (Fig. S3; #4484), two Tpk1-3^as^ cultures were grown to day 4 and each split into six flasks and treated with 0, 1, 10, 50, 100, 500 and 1000 nM 1-NM-PP1 for 15 min. Glucose was added to each culture 5 min before harvest.

To determine the relative contributions of the PKA isoforms to protein phosphorylation (Fig. 4, Fig. S5; #4584), two replicate cultures of Tpk1-3^as^, Tpk1/2^as^, Tpk1/3^as^ and Tpk2/3^as^ were grown in SG to day 4. Each culture was then split into four flasks. The cells were then harvested (i) during steady state in SG medium, (ii) 5 min after the addition of glucose to the medium, (iii) after 15 min incubation with 500 nM 1-NM-PP1, and (iv) after 10 min pretreatment with 500 nM 1-NM-PP1 followed by 5 min in glucose.

To measure the dephosphorylation rates of the PKA substrates (Fig. 6; #4489), two Tpk1-3^as^ cultures were grown to day 4. The cultures were then treated with glucose for 5 min, followed by 500nM 1-NM-PP1, and samples harvested after 0, 1, 2.5, 5, 10, 15, 30 and 60 min. In parallel, two cultures of wt strain were grown to day 4 and each split into two flasks. Glucose was added for 5 min, before addition of DMSO or 500 nM 1-NM-PP1 for 5 min.

### Phospho-proteomics and sample processing

Yeast cells were fixed by mixing 47 ml of culture with 3 ml trichloro-acetic acid and incubating on ice for at least 30 min. The cultures were then centrifuged at 4000 rpm (Sorvall 75006445 rotor) for 5 mins at 6 °C, most of the supernatant discarded, and the remainder of the culture transferred into a 2 ml screw-cap tube. The 2 ml tubes were then spun at 6000 g for 30 s at 4 °C, the supernatant removed, and the pellet washed with 1 ml cold acetone and sonicated for 5 s at 15% power using a Sonic Dismembrator Model 500 sonicator (Fisher Scientific), fitted with a microtip. The samples were then centrifuged again at 6000 g for 30s, the supernatant removed, and the acetone wash procedure repeated. The resulting cell pellet was then dried in a vacuum centrifuge for 15 min (without heating) and stored at −80 °C for up to a month.

Frozen cell pellets were resuspended in 400 μl urea-buffer (8 M urea, 100 mM ammonium bicarbonate (ABC), 5 mM EDTA), acid washed glass beads added to the top of the tube, and additional urea buffer (about 200 μl) added to create a uniform slurry. The cells were then lysed by bead beating for 1 min at 6500 rpm and 4 °C, and the lysate was spun out of the beads by puncturing the base of the tube with a 23G hot needle and centrifuging at 3000 rpm (Sorvall 75006445 rotor) for 5 min. The final protein concentration in each extract was then measured using a BCA assay (Pierce).

Lysate containing 200 μg of protein was diluted to 1 μg/μl in urea buffer before Tris(2-carboxyethyl)phosphine hydrochloride and iodoacetamide were added to a final concentration of 5 mM each, and the samples incubated for 30 min at room temperature in the dark. The urea concentration was then diluted to 5.5 M with 50 mM ABC and the samples digested with LysC (New England Biolabs) at 1:100 mass-ratio at 37°C for 3 hours with agitation. The urea concentration was then diluted to 1 M with 50 mM ABC and the proteins digested with trypsin (Promega, V511c) at 1:100 mass-ratio at 37 °C overnight.

The samples were then acidified by adding 20% TFA to a final concentration of 1% v/v, clarified by centrifugation (21,000 g), and desalted on SepPak Plus C18 cartridges (Waters: WAT020515). In the desalting step, cartridges were first flushed with 5 ml solution B (65% acetonitrile (MeCN), 0.1% trifluoroacetic acid (TFA)) and 10 ml solution A (2% MeCN, 0.1% TFA) using a vacuum manifold. The peptides were then diluted in 8 ml solution A and run into the cartridges, flushed with 10 ml solution A, and eluted 2 times with 600 μl of solution B. The peptides were then dried in a vacuum centrifuge and stored at −80 °C for up to 3 weeks.

Samples were enriched for phosphopeptides using magnetic Ti(IV)-IMAC beads (MagReSyn MR-TIM005) following the manufacturer’s instructions. In brief, peptides were resuspended in 200 μl loading solution (0.1M glycolic acid in 80% MeCN, 5% TFA) and incubated with 40 μl pre-washed beads. The beads were then washed with loading solution, 80% ACN, 1% TFA and 10% ACN, 0.2% TFA and the phosphopeptides eluted in 2x 135 μl 1% NH4OH into 90 μl 10% formic acid (FA).

Finally, the phosphopeptides were desalted on micro-spin columns (Nest Group: SEM SS18V) with centrifugation at 1500 g for 1 minute at each step: Columns were conditioned with 400 μl 90% MeCN, 0.1% FA and equilibrated with 350 μl 5% MeCN, 0.1% FA. Peptides were applied, columns washed with 350 μl 5% MeCN, 0.1% FA and peptides eluted with 200 ul 50% MeCN, 0.1% FA and dried in a vacuum centrifuge.

### Mass Spectrometry

Samples were resuspended in 10.5 μl 0.1% FA, of which 1.5 μl were injected for HPLC-ESI-MS/MS analysis. Data acquisition was performed in positive ion mode on a Thermo Scientific Orbitrap Fusion Lumos tribrid mass spectrometer fitted with an EASY-Spray Source (Thermo Scientific, San Jose, CA). NanoLC was performed using a Thermo Scientific UltiMate 3000 RSLCnano System with an EASY Spray C18 LC column (Thermo Scientific, 50 cm x 75 μm inner diameter, packed with PepMap RSLC C18 material, 2 μm, cat. # ES803): loading phase for 15 min at 0.300 μl/min; linear gradient of 1–34% Buffer B in 119 min at 0.220 μl/min, followed by a step to 95% Buffer B over 4 min at 0.220 μl/min, hold 5 min at 0.250 μl/min, and then a step to 1% Buffer B over 5 min at 0.250 μl/min and a final hold for 10 min (total run 159 min); Buffer A = 0.1% FA; Buffer B = 0.1% FA in 80% ACN. Spectra were collected using XCalibur, version 2.3 (ThermoFisher Scientific). Precursor scans were acquired in the Orbitrap at 120 000 resolution on a mass range from 375 to 1575 Th. Precursors were isolated with an isolation width of 1.6 Th and subjected to higher energy collisional dissociation. MS/MS scans were acquired in the ion trap on the m/z range of 120 to 2000 Th with a fill time of 35 ms.

### Phospho-proteomics data analysis

Progenesis QI for proteomics software (version 2.4, Nonlinear Dynamics Ltd., Newcastle upon Tyne, UK) was used to perform ion-intensity based label-free quantification as described previously (Parker et al., 2019). In brief, in an automated format, .raw files were imported and converted into two-dimensional maps (y-axis = time, x-axis =m/z) followed by selection of a reference run for alignment purposes. An aggregate data set containing all peak information from all of the samples in a given experiment was created from the aligned runs, which was then further narrowed down by selecting only +2, +3, and +4 charged ions for further analysis. A peak list of fragment ion spectra was exported in Mascot generic file (.mgf) format and searched against a UniProt *S. cerevisiae* S288c database (6728 entries) using Mascot (Matrix Science, London, UK; version 2.6). The search variables that were used were: 10 ppm mass tolerance for precursor ion masses and 0.5 Da for product ion masses; digestion with trypsin; a maximum of two missed tryptic cleavages; variable modifications of oxidation of methionine and phosphorylation of serine, threonine, and tyrosine; 13C=1. The resulting Mascot .xml file was then imported into Progenesis, allowing for peptide/protein assignment, while peptides with a Mascot Ion Score of <25 were not considered for further analysis.

Peptide ion data were exported as a .csv file. Positions of phosphorylation sites within the protein were obtained by mapping the peptide sequence to the protein sequence in the .fasta file used for the database search. Duplicate entries of the same peptide ion mapping to more than one protein were collapsed to one entry. Normalized intensities of singly phosphorylated peptides mapping to the same phosphosite were averaged, while data from multiply-phosphorylated peptides was dropped from the analysis.

To calculate fold-changes, flooring was implemented by adding 1000 counts to all normalized intensities (at a median normalized intensity around 4× 10^4^). Significance of differences between sample means were evaluated using a two-sided Student t-test. False discovery rate (FDR) calculations were performed on scrambled data, where normalized intensities for each phosphosite were randomly assigned to samples. The FDR was calculated by dividing the number of phosphosites passing the applied threshold by the number from the original dataset and averaging over ten scrambled datasets.

To generate sequence logos, sequence windows of +/− 5 amino acids around each phosphosite were extracted. Sequences corresponding to phosphosites determined as PKA-dependent were compared to all quantified sites using pLogo (O’Shea et al., 2013). The significance of enrichment of sub-motifs RRX<u>S</u>, RKX<u>S</u>, RZX<u>S</u>, KRX<u>S</u>, ZRX<u>S</u>, KKX<u>S</u>, KZX<u>S</u>, ZKX<u>S</u> and [R/K][R/K]X<u>T</u>, where the underlined residue is the phosphosite, X is any amino acid and Z is any amino acid except K or R, was calculated from a binomial distribution using the frequency of a motif among all quantified sites as its baseline probability.

### Proliferation assays

For proliferation assays in glucose, three replicate cultures of the Tpk1-3^as^ and the base wild-type strain were inoculated in SG medium on day 0 and diluted in the same medium on day 2 and day 3. Each culture was then split into nine flasks and three treated 100 nM 1-NM-PP1, three treated with 500 nM 1-NM-PP1, and three treated with an equivalent volume of DMSO. 2% glucose was then added to each flask (from a 40% stock) after 15 min. Two additional control cultures were also grown where water was added instead of glucose to track the base growth rate. Samples were taken at multiple time-points and ODs determined using a spectrophotometer (GENESYS 30; Thermo Scientific) with a 1 cm light path. Cultures were diluted in pre-warmed medium with the appropriate amount of glucose and 1-NM-PP1 once they reached OD 0.8-1.0. These dilution factors are factored into the reported ODs. Proliferation rates in SG were determined in an analogous manner, but glucose was never added to the flasks.

## ACKNOWLEDGEMENTS

We thank Paul Langlais and Austin Lipinski for mass spectrometry data acquisition, Pascale Charest, Arminja Kettenbach, Telsa Mittermeier, Andrew Paek and the members of the Capaldi lab for helpful advice and Carol Dieckmann for critical reading of the manuscript. We acknowledge financial support through a BIO5 Postdoctoral Fellowship (MJP), the UBRP program including funding from Bio5 and the Office of Research Innovation and Discovery (NC and BM), the Western Alliance to Expand Student Opportunities (WAESO) Louis Stokes Alliance for Minority Participation (LSAMP) National Science Foundation (NSF) Cooperative Agreement No. HRD-1101728 (NC), and NIH 1R01GM097329 (APC).

## AUTHOR CONTRIBUTIONS

M.P. and A.C. conceived the study. M.P. performed most experiments and analyzed the results with A.C. N.C. performed and analyzed proliferation assays. B.M. contributed to proteomics sample collection. M.P. and A.C. wrote the manuscript.

---

**Fig. S1.**
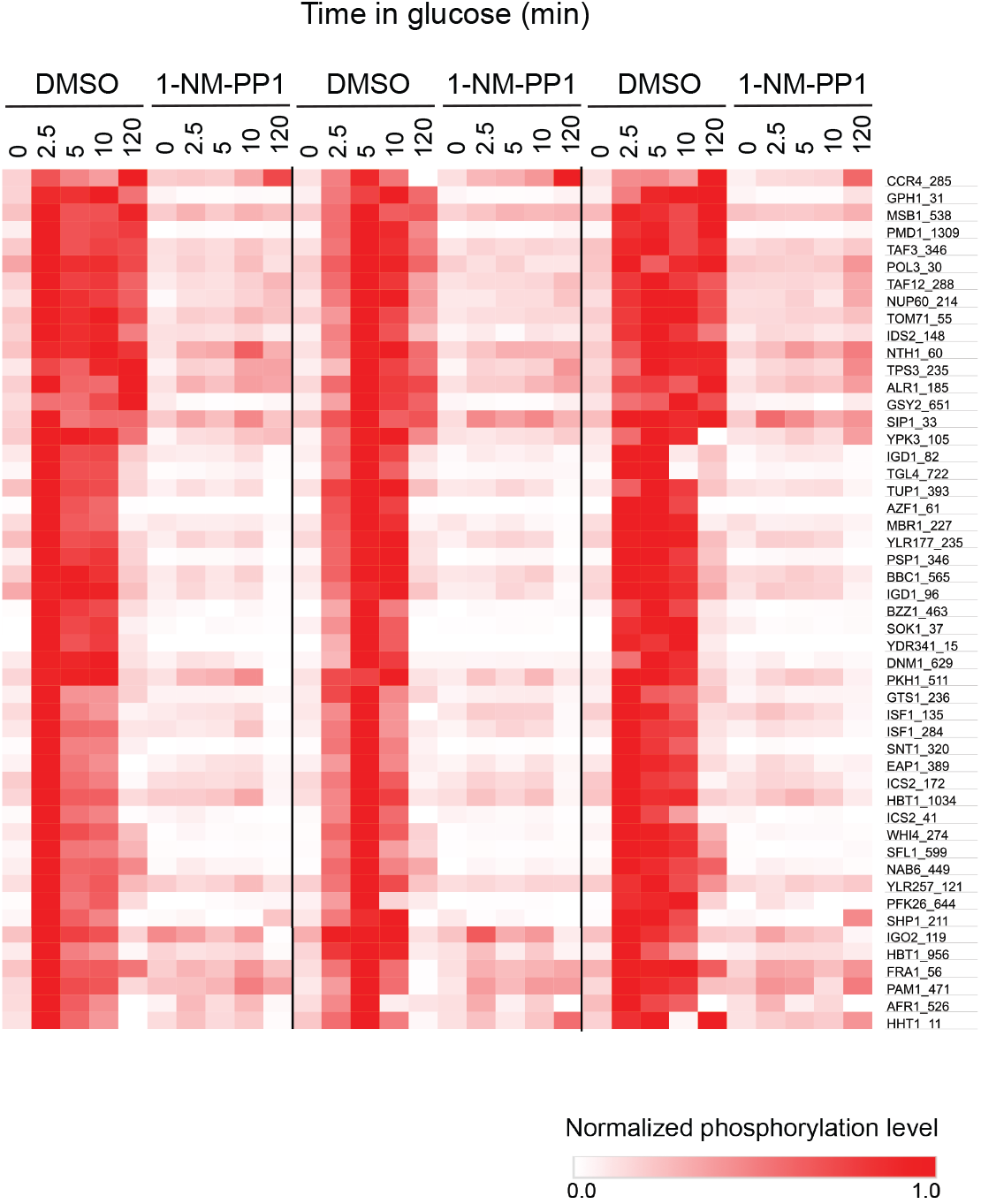
Reproducibility of the time-course data. Heatmap showing the phosphorylation level of the PKA-dependent sites as a function of time across three biological replicates, each normalized to the 5 min timepoint w/o 1-NM-PP1. The data are the same as in Fig. 1b but are shown without averaging over replicates.

**Fig. S2.**
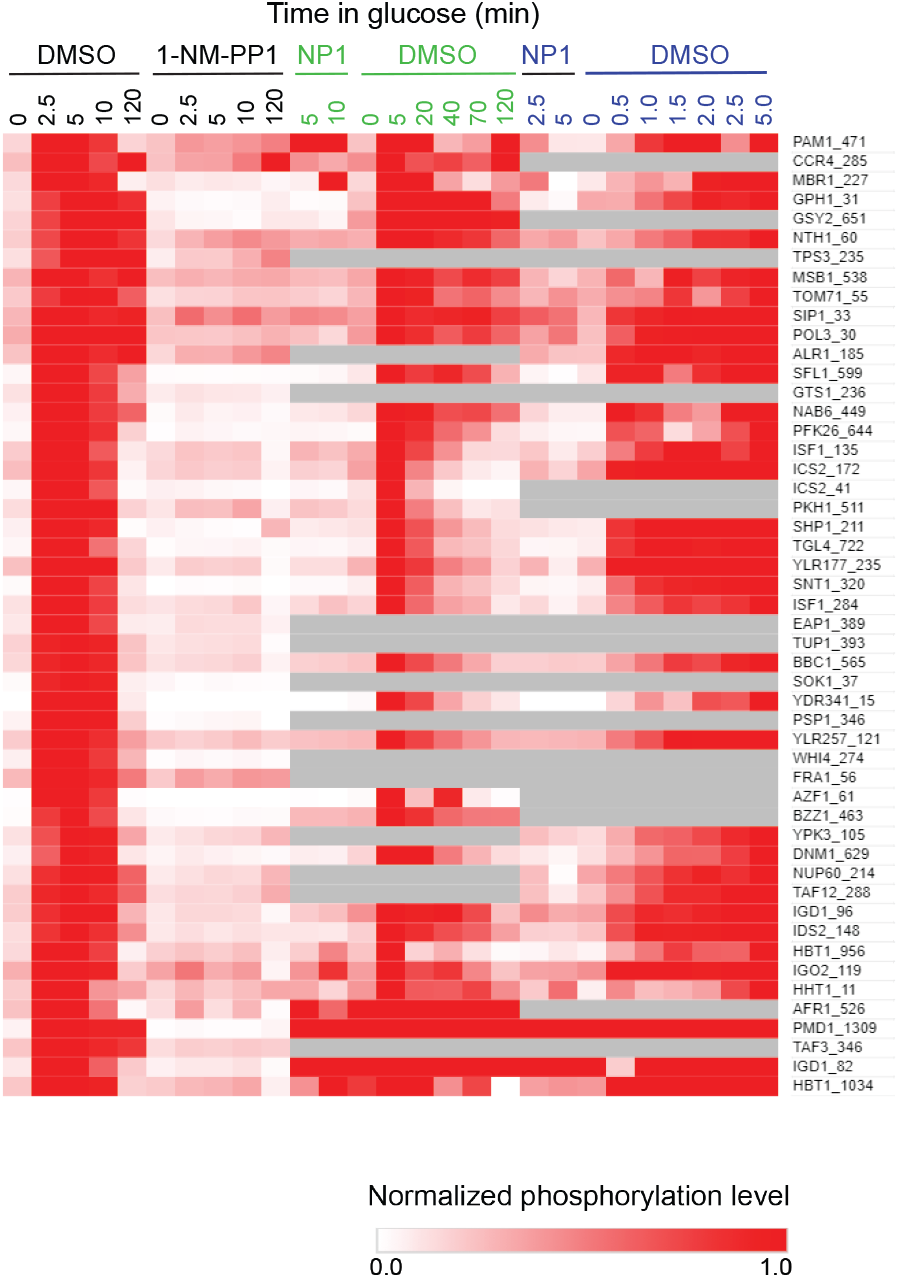
Extended time-course data. Heatmap showing the average phosphorylation level of the PKA-dependent sites at early and intermediate timepoints, along with the data from Fig. 1. Experiments were carried out identically to those in Fig. 1, except that the cells were harvested a different time-points. All values are normalized to the data from the 5 min timepoint w/o 1-NM-PP1 from the matching experiment.

**Fig. S3.**
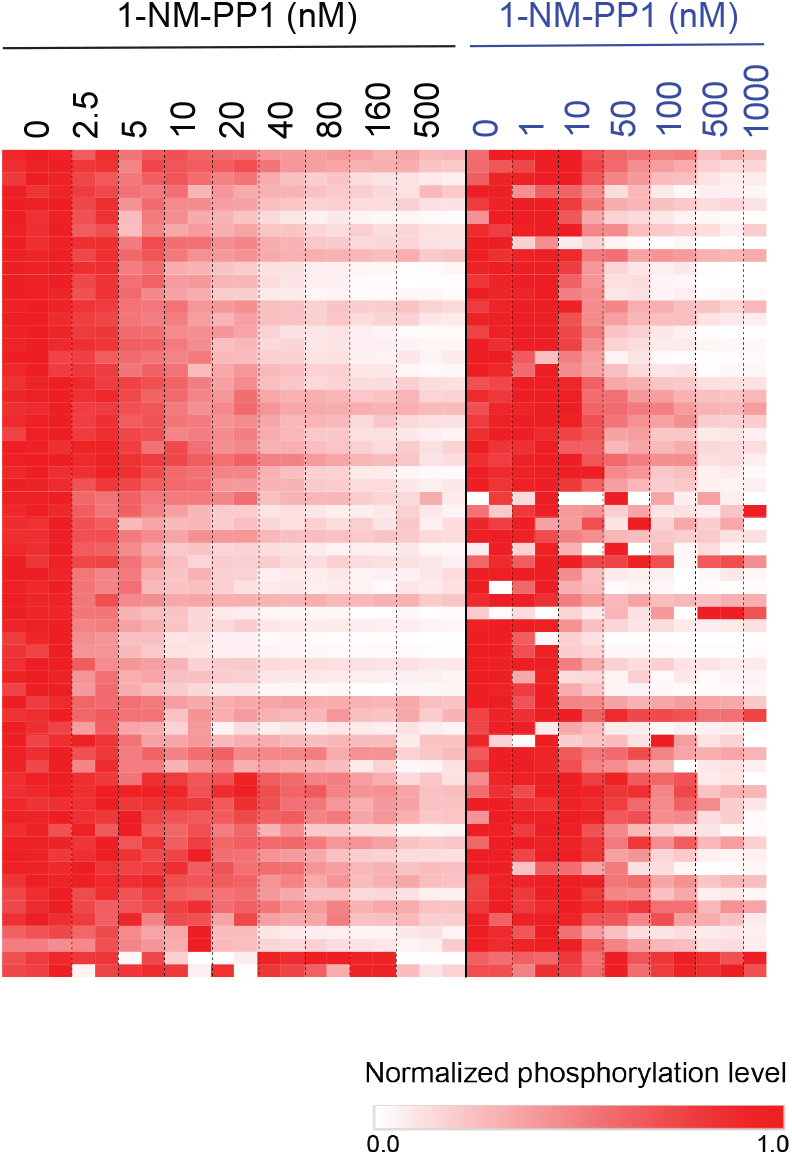
Reproducibility of the 1-NM-PP1 sensitivity data. Tpk1-3^as^ cells were grown in medium with glycerol as the carbon source, pre-treated with the indicated concentrations of 1-NM-PP1 for 10 min, and then exposed to glucose for 5 min and analyzed using phosphoproteomics. The signal at each phosphosite is normalized to its maximum value in the matching experiment. The left side of the heatmap (black labels) shows the data from the experiment used to calculate the average values in Fig. 3. The right side of the heat map (blue labels) shows the data from a separate experiment used to further confirm the reproducibility 1-NM-PP1 sensitivity plots.

**See attached Pdfs**

**Fig. S4**. 1-NM-PP1 dose-response curves for all 83 phosphosites that are classified as PKA-dependent and contain at least one arginine or two lysines in the P-3 and P-2 positions. The red curves show the best fit to the a standard binding equation (K_i_/(K_i_ +[NM-PP1]) + c, where K_i_ is the inhibition constant and c is the baseline level of phosphorylation). The dashed violet lines show the best fit value of c. (**a, b**) panels showing the sites that fit well (K_i_ with a relative SE<0.33), and poorly to the binding equation, respectively.

**Fig. S5.**
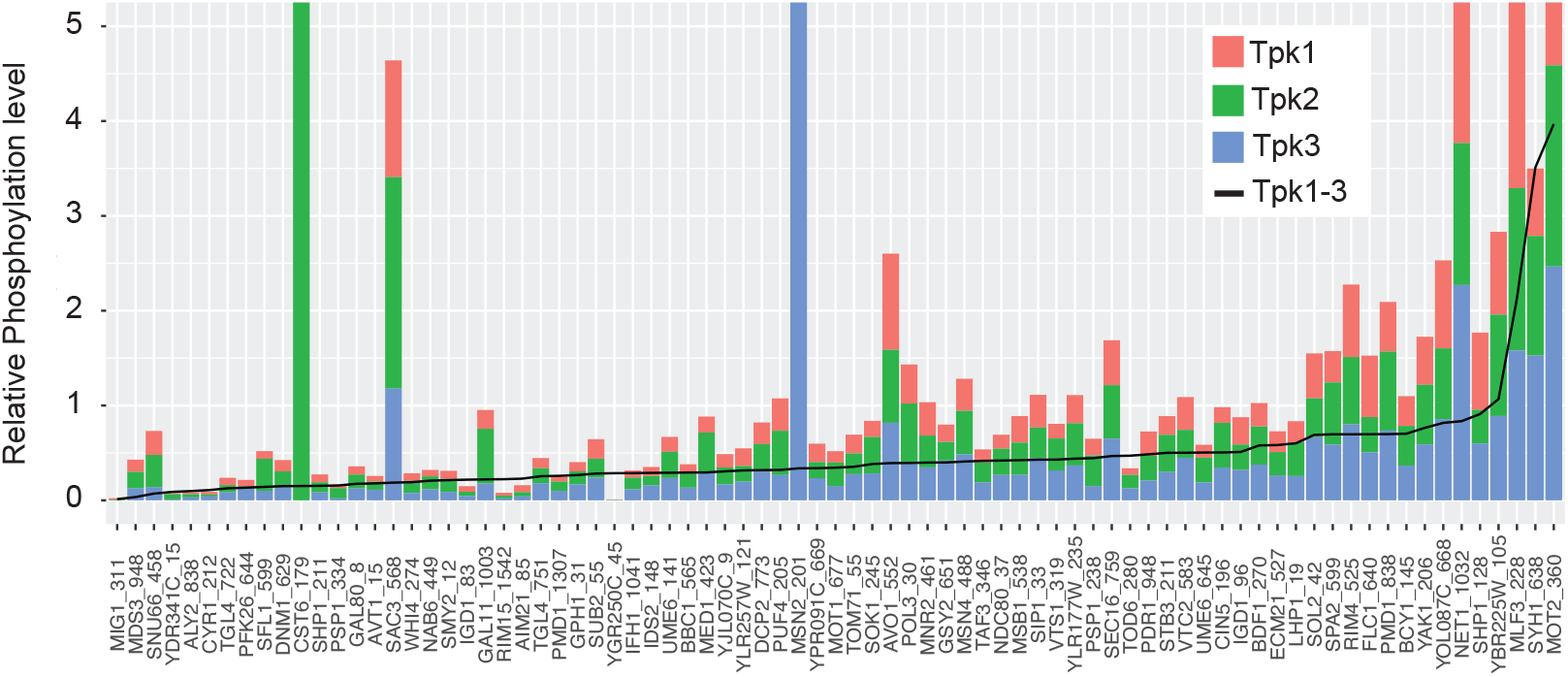
Influence of the three Tpk isoforms on protein phosphorylation. The stacked bar plot shows the phosphorylation level at PKA-dependent phosphosites in SG when a single Tpk isoform is active. The black line shows the phosphorylation level when all three Tpk isoforms are active (the data are from the same experiment as shown in Fig. 4a).

**Fig. S6.**
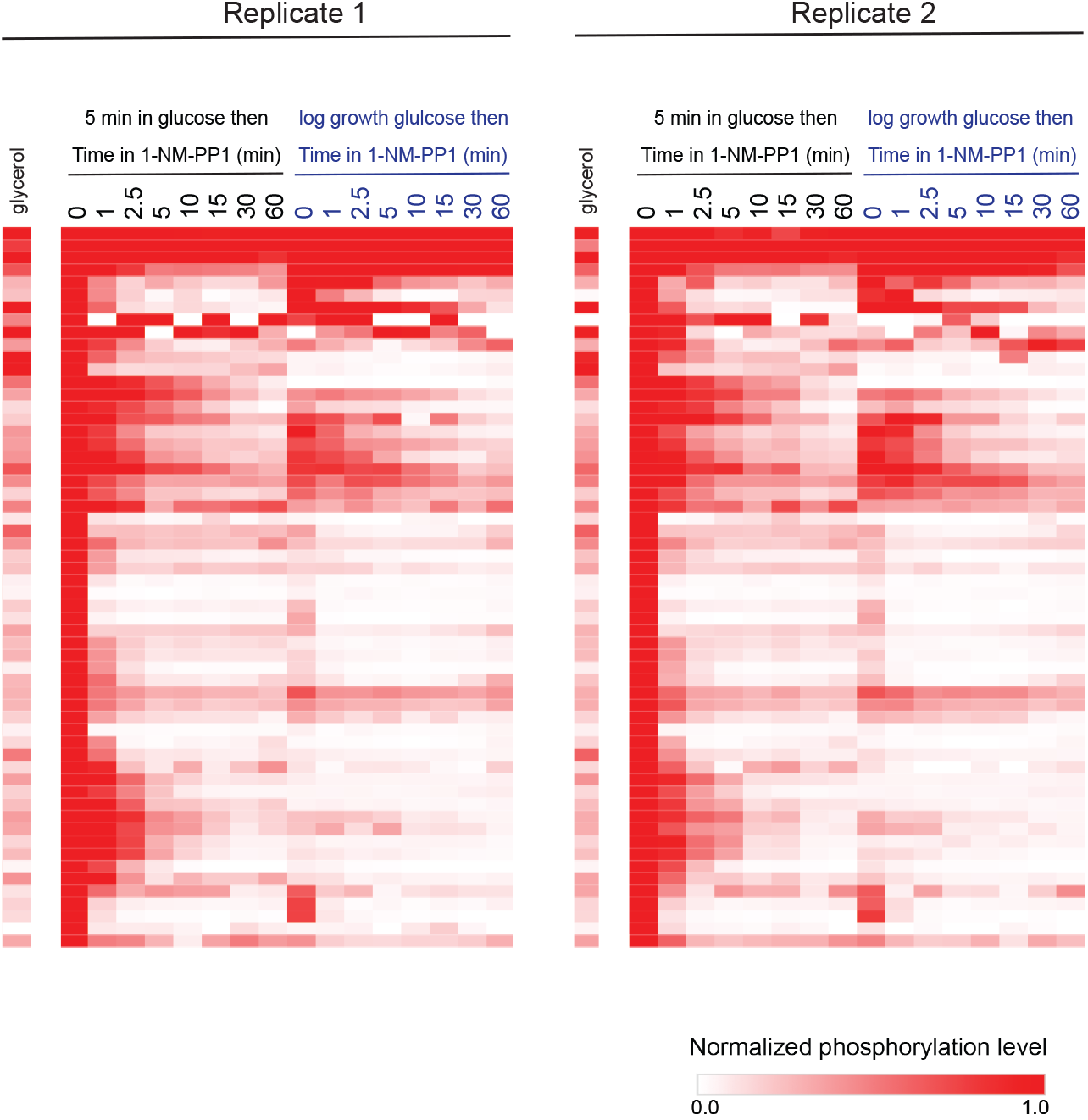
PKA-dependent phosphorylation decreases rapidly after PKA inhibition. In one experiment (carried out in duplicate), Tpk1-3^as^ cultures were grown in SG medium and incubated with glucose for 5 min (defined as t = 0), 500 nM 1-NM-PP1 was then added, and the phosphorylation levels followed over time. In a second experiment (carried out in duplicate), cultures were grown in synthetic medium with glucose, 500 nM 1-NM-PP1 added at t = 0, and phosphorylation monitored over time. The heatmap shows the data for all PKA-dependent phosphosites that were detected, and each value is normalized to that at time 0 in the matching experiment.

**See attached pdfs**

**Fig. S7**. Extended 1-NM-PP1 dependent dephosphorylation data. Complete set of dephosphorylation curves from Fig. 6. (**a, b**) panels showing the sites that fit well (with a relative SE<0.33) and poorly to a single exponential equation, respectively.

**See attached excel spreadsheet**

**Table S1.** Raw data for all phosphoproteomic experiments carried out in this study. The experiments shown in each figure are split into different spreadsheets, as labelled on each tab.

