## Supplementary figures and images for "Systems level analysis of time and stimuli specific signaling through PKA"

### Fig. S4a

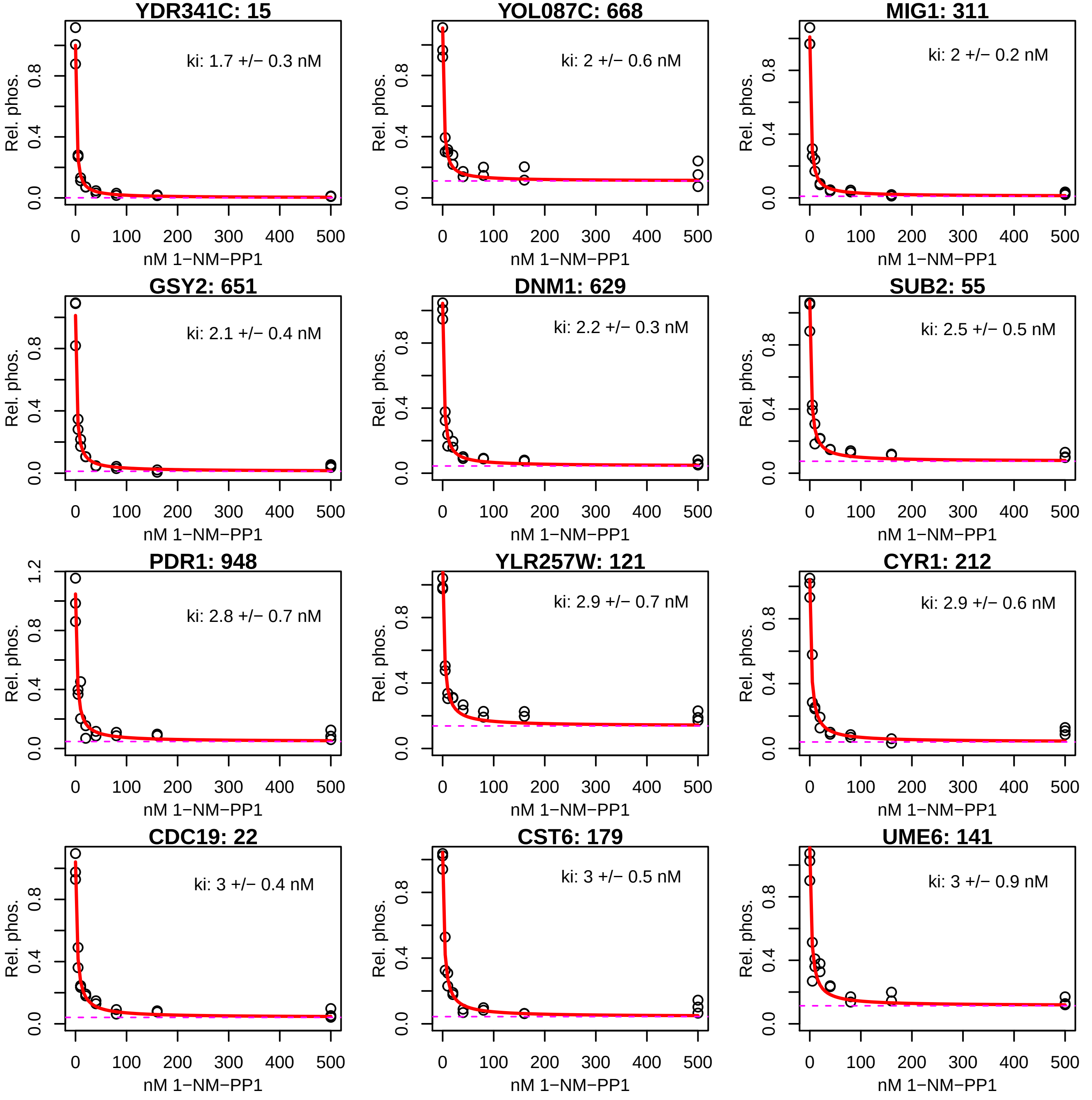

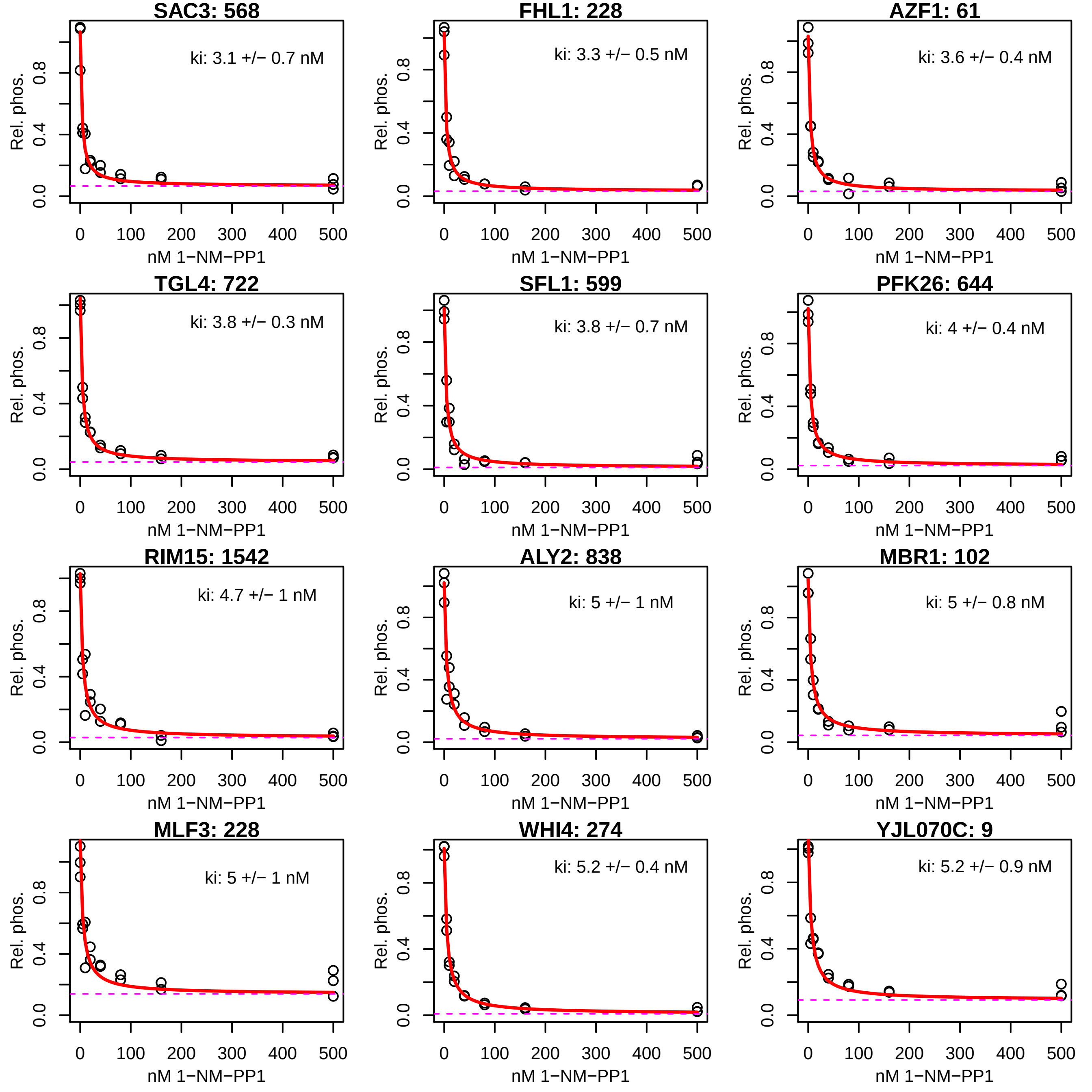

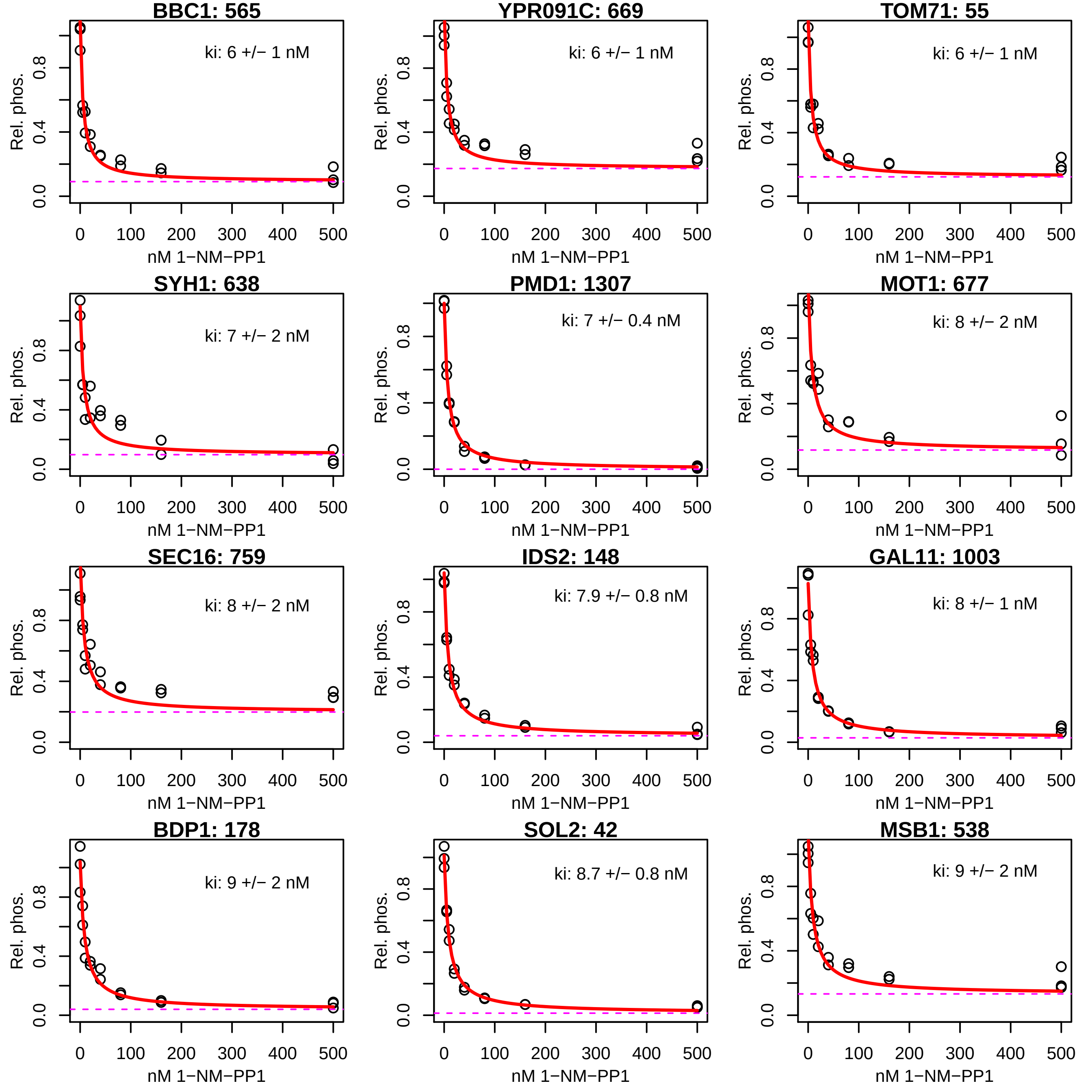

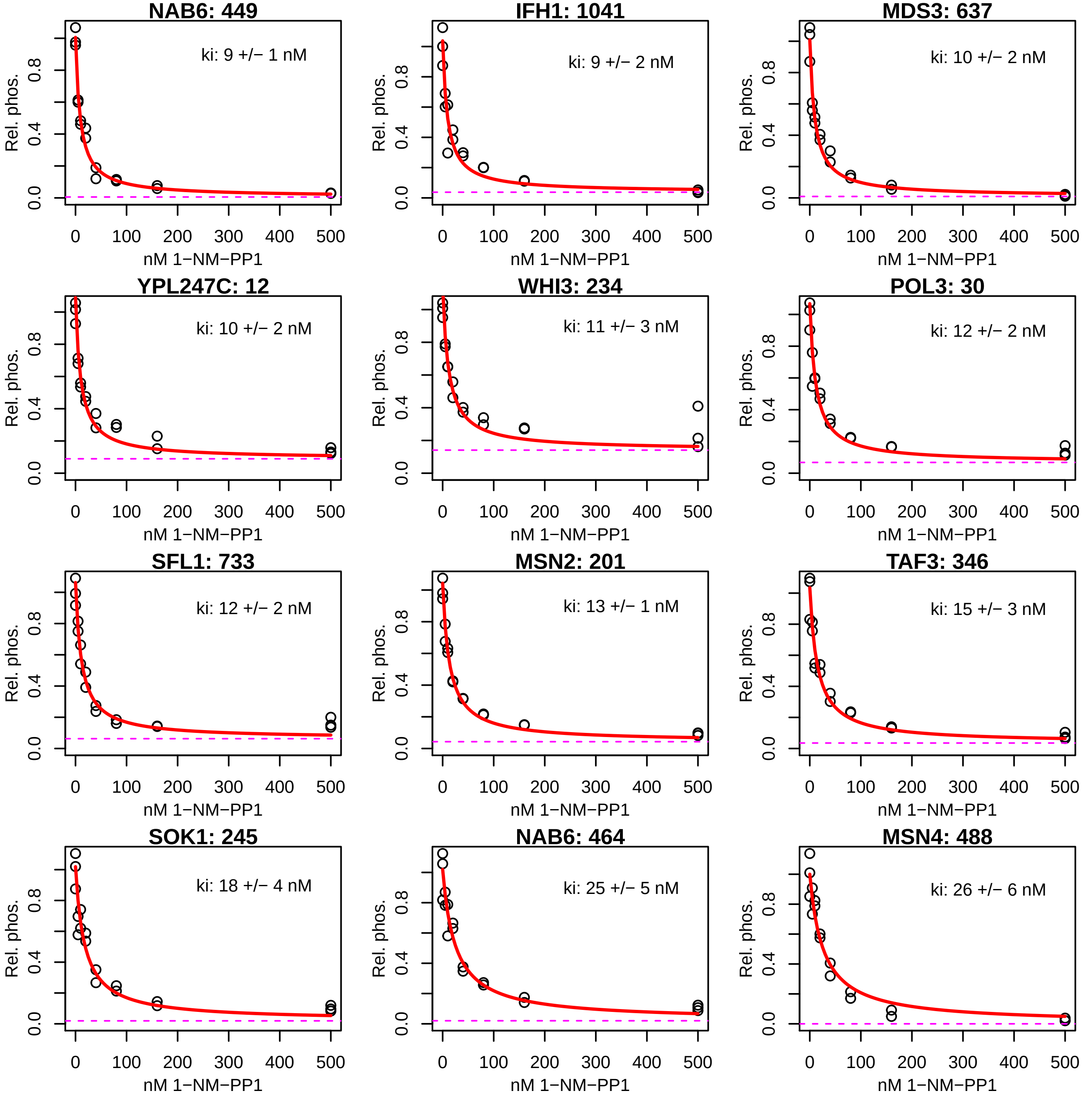

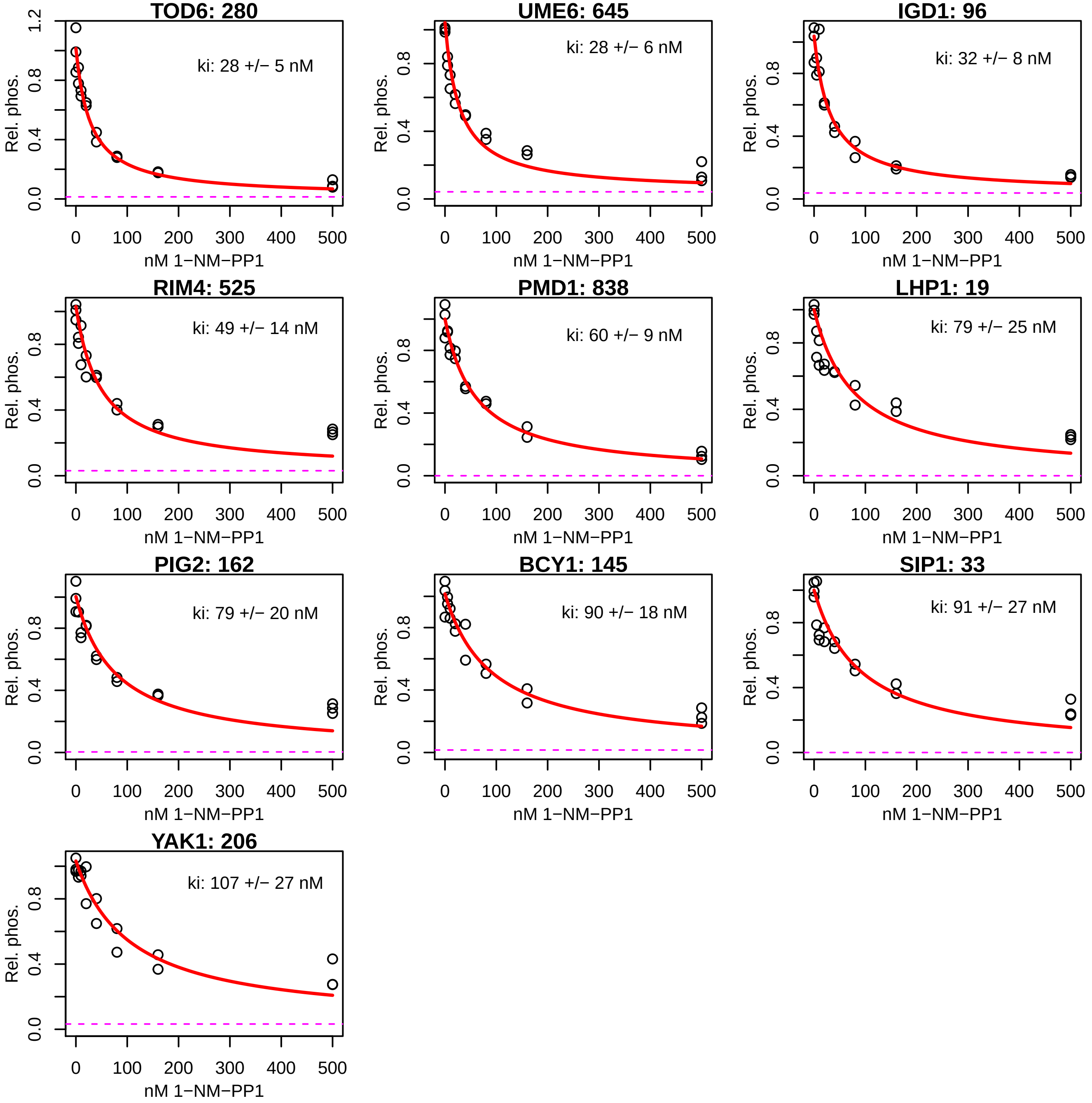

### FIg. S4b

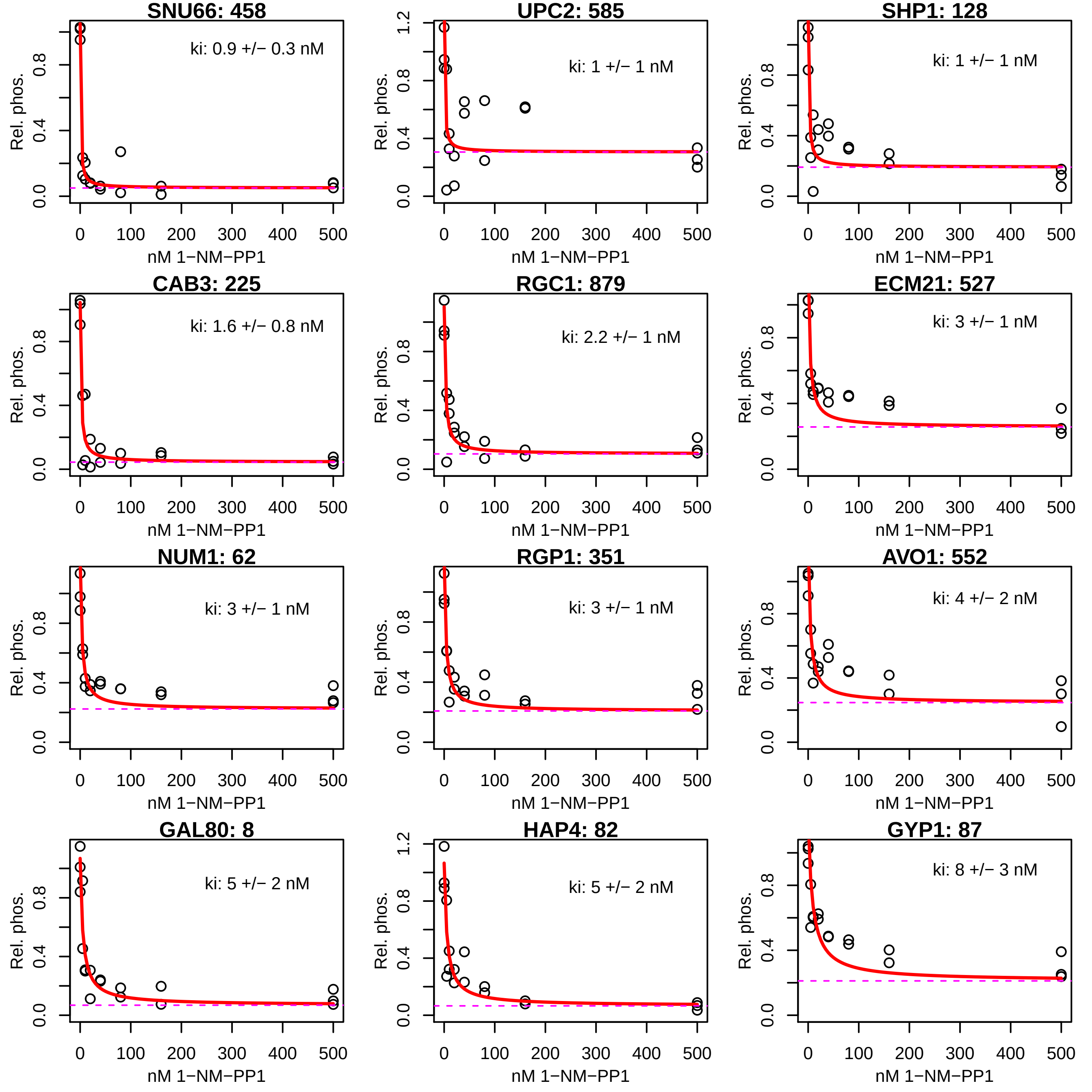

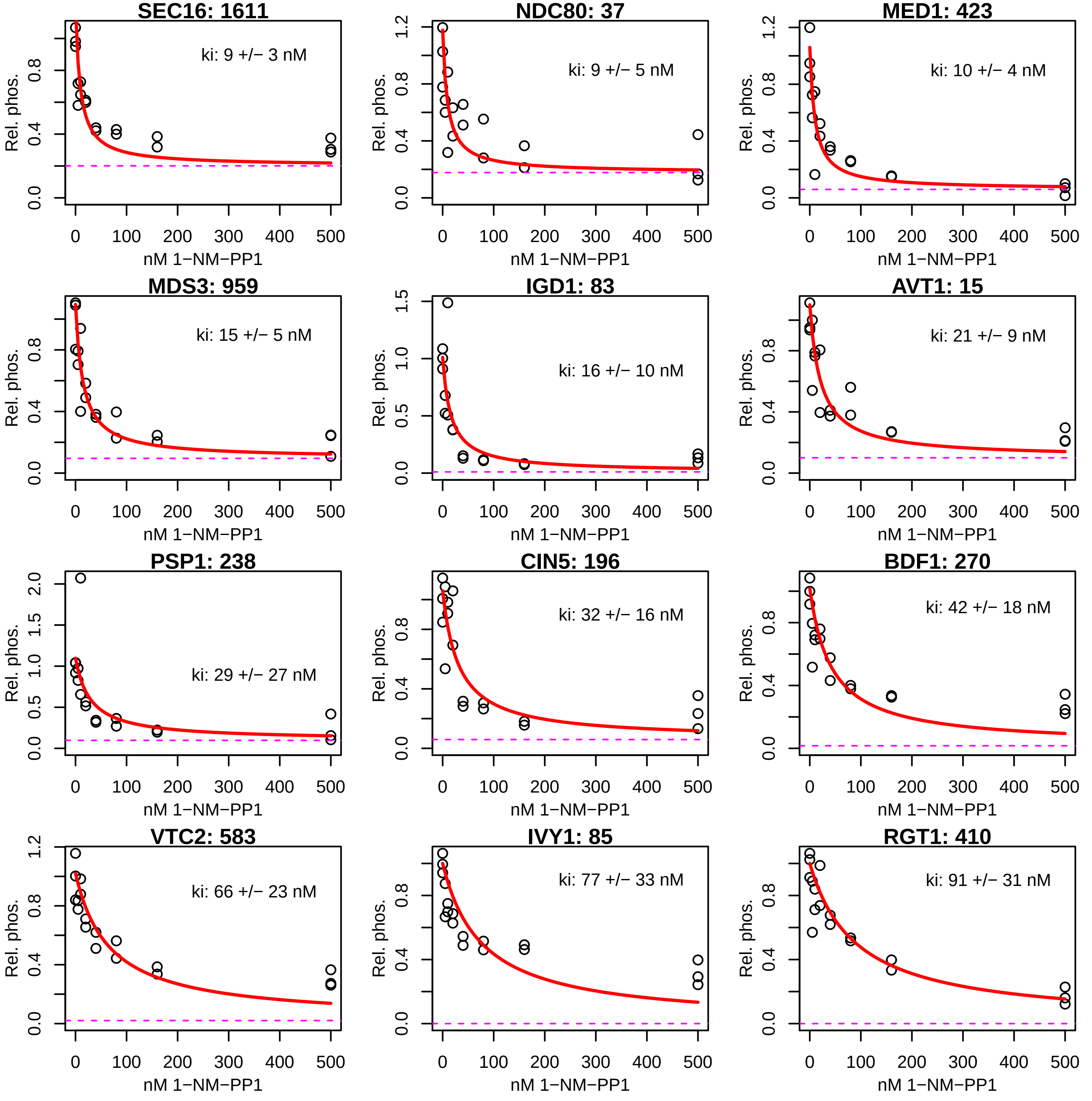

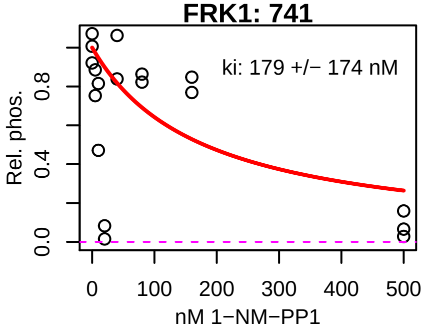

### FIg. S7a

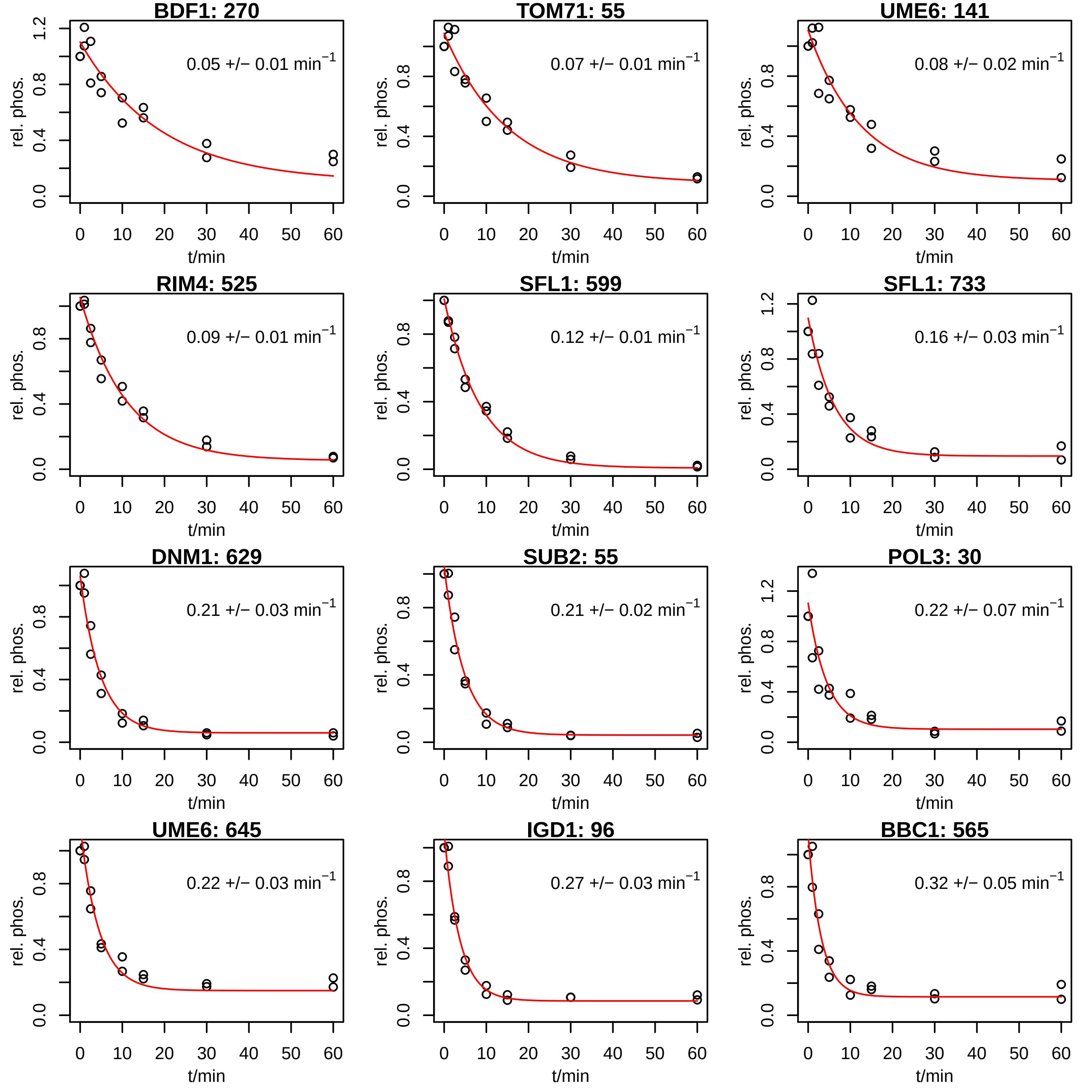

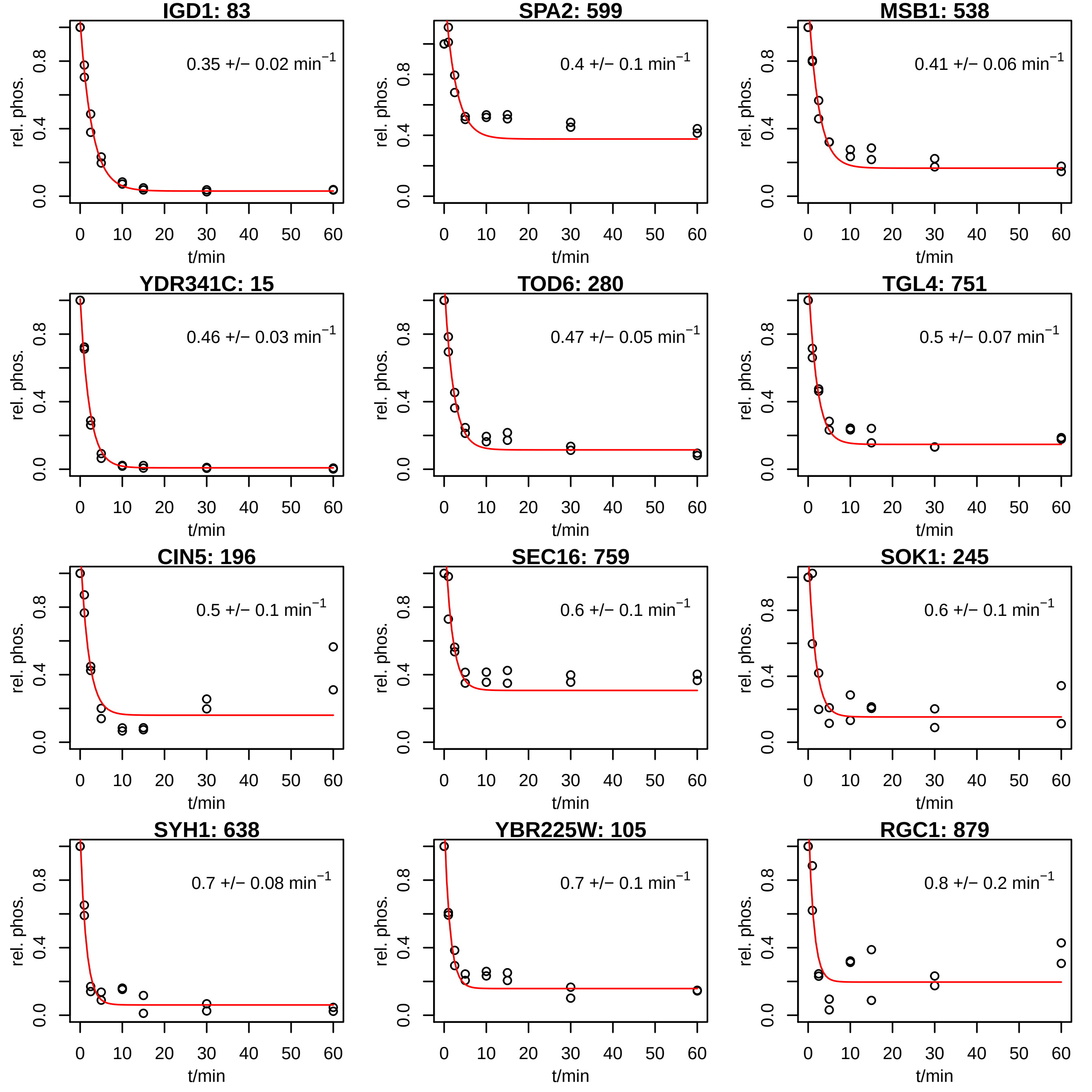

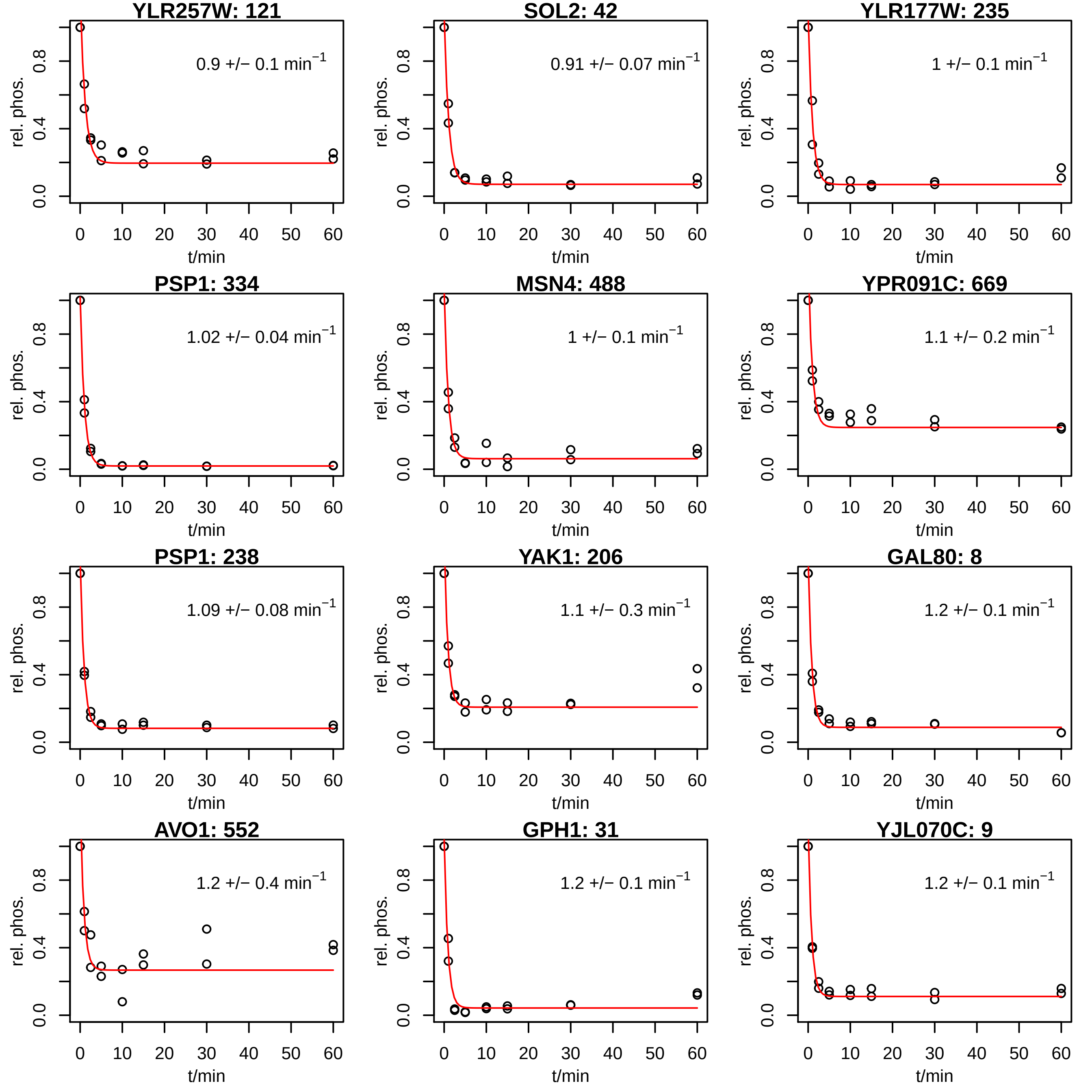

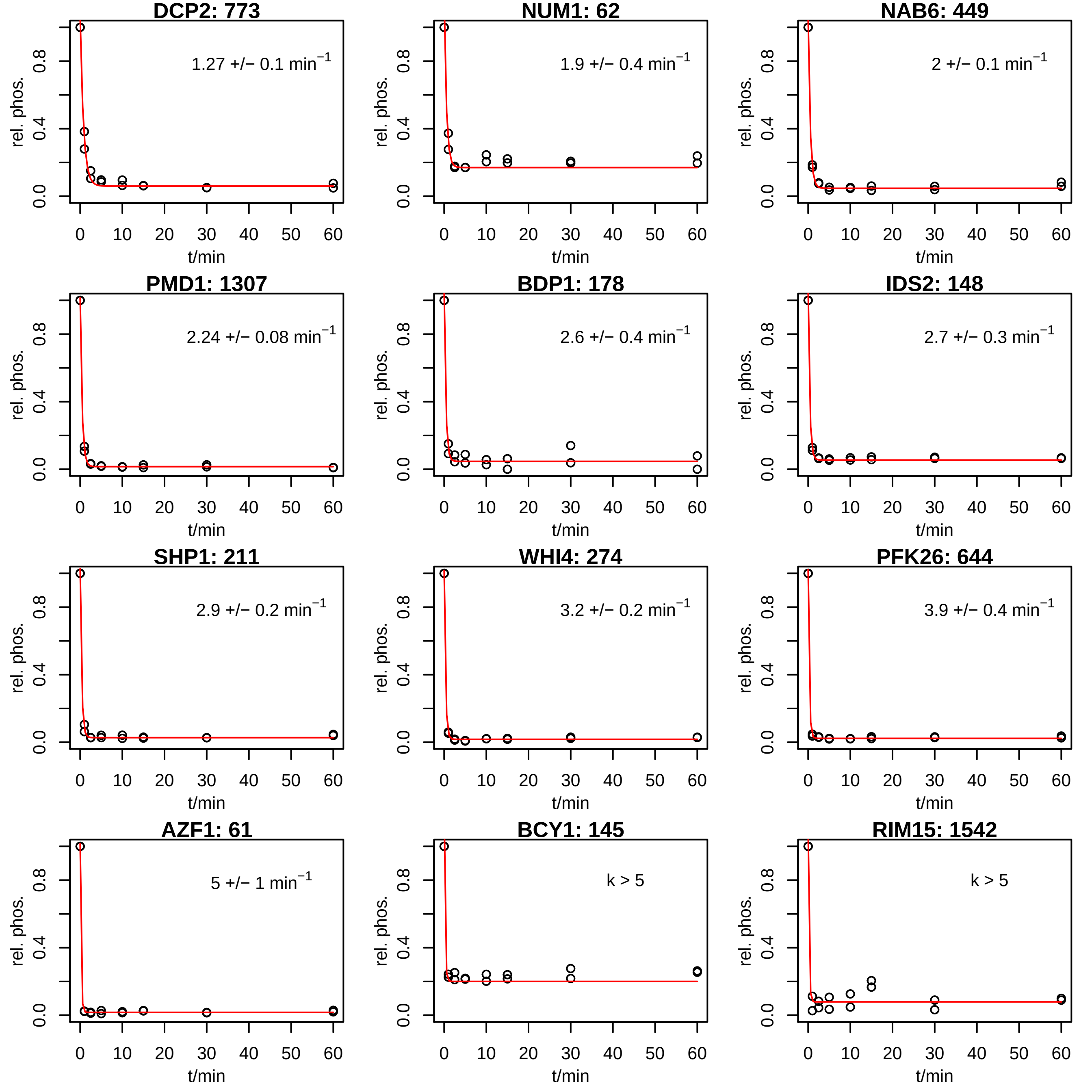

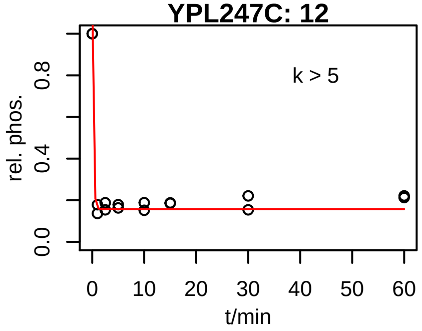

### FIg. S7b

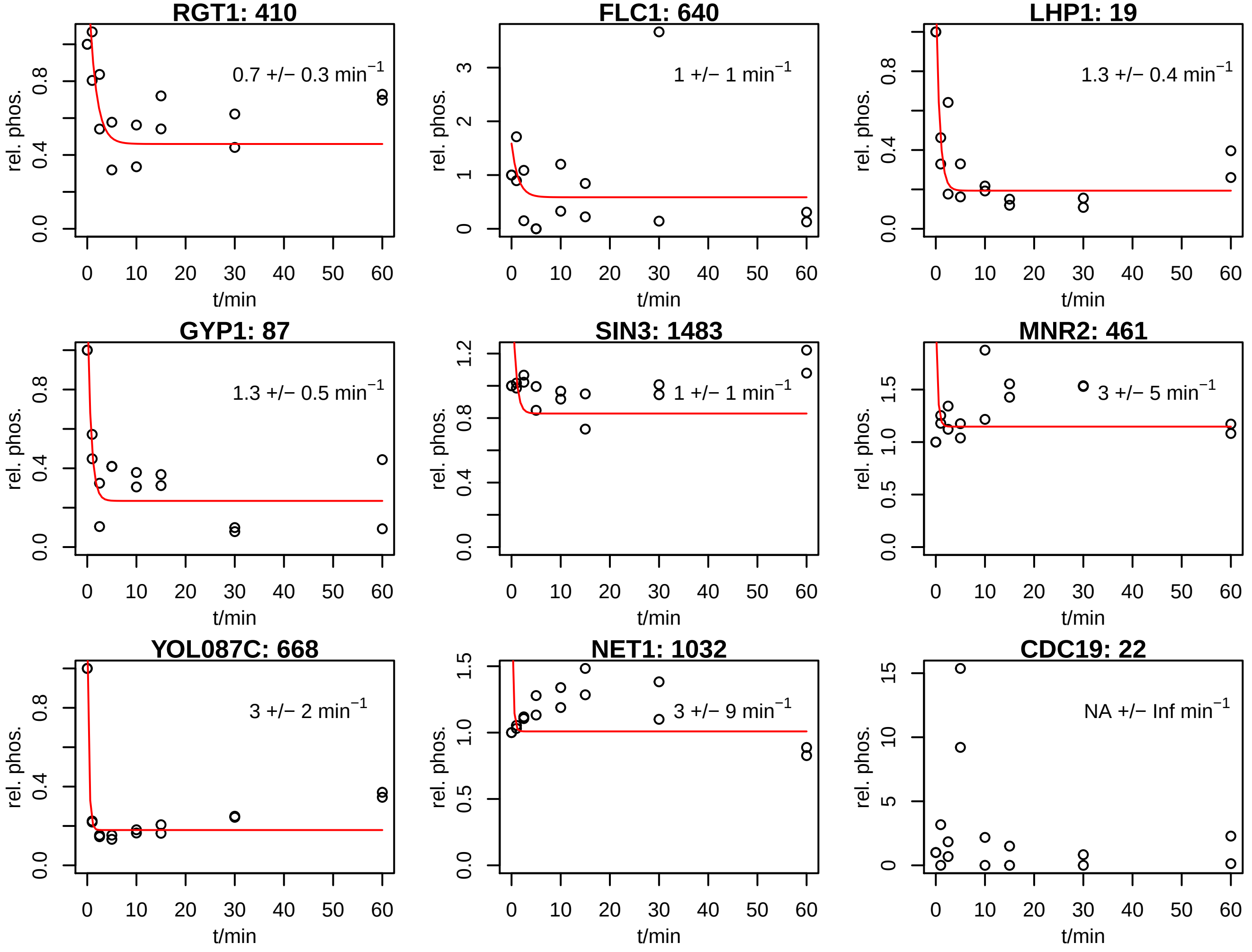
